# White matter and literacy: a dynamic system in flux

**DOI:** 10.1101/2022.06.21.497048

**Authors:** Ethan Roy, Adam Richie-Halford, John Kruper, Manjari Narayan, David Bloom, Pierre Nedelec, Leo P. Sugrue, Andreas Rauschecker, Timothy T. Brown, Terry L. Jernigan, Bruce D. McCandliss, Ariel Rokem, Jason D. Yeatman

**Author notes:** Corresponding Author, Graduate School of Education, Stanford University, 482 Galvez Mall, Stanford, CA, 94305, USA.

## Abstract

Cross-sectional studies have linked differences in white matter tissue properties to reading skills. However, past studies have reported a range of, sometimes conflicting, results. Some studies suggest that white matter properties act as individual-level traits predictive of reading skill, whereas others suggest that reading skill and white matter develop as a function of an individual’s educational experience. In the present study, we tested two hypotheses: a) that diffusion properties of the white matter reflect stable brain characteristics that relate to reading skills over development or b) that white matter is a dynamic system, linked with learning over time. To answer these questions, we examined the relationship between white matter and reading in a five-year longitudinal dataset and a series of large-scale, single-observation, cross-sectional datasets (N=14,249 total participants). We find that gains in reading skill correspond to longitudinal changes in the white matter. However, in the single-observation datasets, we find no evidence for the hypothesis that individual differences in white matter predict reading skill. These findings highlight the link between dynamic processes in the white matter and learning.

## 1. Introduction

White matter - the tissue that contains the long-range axonal connections among different brain regions - was historically viewed as static infrastructure critical for healthy cognitive function(1, 2). However, the field went through a sea change as it became clear that oligodendrocytes, the glial cells responsible for myelination in the white matter, actively monitor neural activity(3, 4). Through signaling mechanisms that sense neuronal discharges, oligodendrocytes actively change properties of the white matter in response to fluctuations in neural activity(5, 6). Indeed, it is now largely appreciated that the white matter not only plays an essential role in behavior, but also that plasticity in the white matter is a critical component of the learning process(4).

Given that white matter is sculpted by experience, and that properties of the white matter can undergo dramatic changes over timescales ranging from hours(7, 8), to days(9), to years(10, 11), we now must grapple with how to interpret individual differences in white matter structure. It is often assumed that white matter differences are static traits that explain differences in behavior and can serve as useful biomarkers of, for example, learning disabilities(12–15). This perspective is supported by dozens of studies that have: a) observed correlations between white matter diffusion properties and behavioral measures of academic skills or b) found group differences in white matter diffusion properties between individuals with versus without learning disabilities(16–18).

Reading serves as a paradigmatic example of the relationship between white matter and academic abilities. Seminal research by Klingberg and colleagues(19) discovered differences in white matter diffusion properties between dyslexic and control subjects, inspiring dozens of labs around the world to try to characterize the white matter phenotype of dyslexia (and reading abilities more broadly)(18, 20–24). Although much of this work has identified white matter differences between strong and struggling readers, these studies have reported a variety of, and sometimes even conflicting, results (see Supplementary Table 1 for an overview of past findings). These incongruencies also appear in various meta-analyses, with some identifying a link between reading ability and the white matter(17) and others finding no brain-behavior relationship in the domain of reading(25). Regardless of the direction of these effects, many of these studies have interpreted observed group differences in the white matter properties of dyslexic versus control participants as a static brain-behavior relationship that can be used to classify an individual based on intrinsic properties of their brain.

In contrast to studies exploring biomarkers that emphasize static brain-behavior relationships, longitudinal studies have highlighted the dynamic relationship between white matter plasticity and learning(9, 26–28). These studies offer compelling evidence that rates of white matter development differ dramatically among individuals and are influenced by an individual’s learning experiences. For example, Yeatman and colleagues (2012) showed that over a three year period subjects with above-average reading skills showed increases in fractional anisotropy (FA), a quantity measured with diffusion MRI, in specific white matter pathways. Struggling readers, on the other hand, showed declines in FA in those same white matter pathways (27). Similarly, Wang and colleagues (2017) showed divergent developmental trajectories for the core reading circuitry at the very beginning of reading instruction(28). Furthermore, intervention studies have identified dramatic changes in the diffusion properties following intensive reading interventions(29, 30). Even over the course of a couple weeks of intensive intervention, white matter tissue properties can change with large effect sizes, calling into question the stability of cross-sectional group comparisons(30). These studies challenge the assumption that white matter differences observed in poor readers are rooted in static brain traits, and instead suggest that the white matter is a dynamic, experience-dependent system that is linked with learning over development.

Until recently, the vast majority of published brain-behavior relationships have relied on small (n<50), relatively homogenous participant samples, recruited from the surrounding community of a single research institution(17). As highlighted by recent work(31), results from small and homogenous samples often do not generalize to a broader population. Resolving this issue has only recently become possible as the neuroimaging community has come together to produce large, publicly available, multi-site datasets that sample a much broader swath of the population than traditional, single-laboratory studies(32–34).

For example, a recent cross-sectional analysis of the Healthy Brain Network (HBN) dataset(32), representing the largest study of white matter and reading to date, did not reveal any significant differences in FA between struggling and typical readers nor a significant relationship between FA and reading scores(35). These findings not only raise questions about the observed relationships between reading and the white matter in smaller, cross-sectional studies, but also about the impact of sample makeup on these brain-behavior correlations, as the HBN dataset is a much more diverse sample than those found in traditional single-lab studies.

In the present study, we seek to adjudicate between two perspectives surrounding the relationship between white matter and academic skills: a) that individual differences in white matter diffusion properties reflect stable brain traits that relate to academic skills or b) that white matter and learning are dynamically linked over time and change in response to an individual’s educational environment. To do so, we capitalize on a large-scale longitudinal dataset to explore the longitudinal relationship between changes in white matter properties and reading development. These data do not reveal any cross-sectional relationships between reading skill and white matter properties but, rather, show that gains in reading skill and changes in the white matter are linked longitudinally. This finding supports the hypothesis that white matter and learning are part of a dynamic system that changes as a function of educational experience.

We then leverage four additional large-scale public datasets, totaling more than 12,000 children and adults, to examine the cross-sectional correlations between white matter and reading ability. In this sample, which is more than two orders of magnitude larger than most previous studies, we find no support for the hypothesis that white matter features serve as static traits that predict differences in reading ability. Our analysis suggests that previous studies were detecting characteristics of small, demographically homogeneous samples. Together these findings suggest that a child’s environment exerts a dramatic influence on white matter development and reject the notion that reading difficulties are explained by a stable white matter phenotype.

## 2. Materials and Methods

To explore questions surrounding the properties of the white matter networks underlying reading skill, we leveraged five large-scale, publicly available datasets. These datasets included the Pediatric Longitudinal Imaging, Neurocognition, and Genetics (PLING(36)) dataset, the Child Mind Institute’s Healthy Brain Network (HBN) dataset (32), the University of California San Diego’s Pediatric, Imaging, Neurocognition, and Genetics (PING(33)), and the Human Connectome Project’s (HCP) Young Adult dataset(73). All five of these datasets provide both neuroimaging data and a variety of behavioral assessments, including raw and age-standardized reading. It should be noted that the diffusion MRI data for the HBN, ABCD, and HCP-YA samples were analyzed using pyAFQ and the PLING, PING, and ABCD samples were analyzed using AtlasTrack. Although tractometry was performed using different computational pipelines, past studies have shown that these analyses are robust to the details of the methodology (43). Code to reproduce all the results in the notebook is available: https://github.com/earoy/longitudinal_wm.

### 2.1 PLING

The PLING dataset was used to explore the relationship between changes in white matter properties and changes in academic skills over time. This dataset tracked 176 individuals over the course of 5 years, although not all subjects participated at all 5 time points. We excluded all subjects who did not participate in at least 3 time points and those that did not pass quality control, we were left with a sample of 73 individuals. At the time of the first observation, subjects were between 5 and 10 years old.

At each time point, participants completed an MRI scanning session as well as an assortment of behavioral tasks. dMRI data was collected at each time point and analyzed with AtlasTrack. This processed diffusion data contains mean FA and MD values for 37 major white matter tracts. The behavioral measures we used in this analysis included a composite reading score derived from the Test of Word Reading Efficiency (TOWRE) Sight Word Efficiency and Phonetic Decoding Efficiency subtests(74).

To explore the longitudinal dynamics between reading skill and diffusion properties of the white matter, we constructed a series of longitudinal models. Although most participants (81%) participated in consecutive timepoints, some skipped an observation and therefore had missing data. Because these data were missing at random, all of our modeling approaches relied on Full Information Maximum Likelihood to generate estimations for these missing values.

First, we generated a linear-mixed effects model predicting mean-centered TOWRE reading scores from time point, initial age, mean-centered FA at each time point, and overall mean FA as fixed effects, and participants as a random effect. The use of mean-centered FA allowed us to discern year-to-year change in FA from overall FA within each participant. To better understand the time series of FA and reading score change, we used a multi-level vector autoregression model (mlVAR(38, 39)). This model used time series data to test if FA at a given time point predicted reading scores at the subsequent time point or vice versa. Finally, we generated a parallel-process latent growth curve model to understand the relationship between the rate of reading score change and the rate of FA development. We constructed this latent growth curve model using the lavaan package in R(75).

### 2.2 HBN

The HBN dataset consists of diffusion MRI and phenotypic data from over 1,500 participants ranging from 5 to 21 years of age from the greater New York City area. In the HBN sample, neuroimaging data were collected at four different scanner sites, however the present analysis only included data from the three sites that used 3T scanners: Rutgers University Brain Imaging Center, the CitiGroup Cornell Brain Imaging Center, and the CUNY Advanced Science Research Center. The sample analyzed in the present study consisted of 777 subjects who passed quality control and had both neuroimaging data and various phenotypic measures, including reading scores and socioeconomic status. To determine reading, we calculated an age standardized composite score based on the word reading and pseudoword decoding subsections of the Wechsler Individual Achievement Test (WIAT(76)). Individuals were labeled as struggling readers if their composite score was below 85 (one standard deviation from the mean). The use of age standardized scores allowed us to control for age-related gains in reading and to generate reading groups that spanned the age range of the entire sample. The raw diffusion MRI data were preprocessed using qsiprep(41) and quality control was performed on the entire dataset using a neural network trained on ratings generated by a combination of diffusion imaging experts and community scientist on two subsets of the HBN dataset(42). Any subjects who did not meet quality control standards were excluded from the analysis.

Once the diffusion imaging data were preprocessed, pyAFQ was used to calculate tractometry properties(43). Briefly, constrained spherical deconvolution(77), implemented in DIPY(78) was used to estimate fiber orientation distributions in every voxel, and probabilistic tractography was used to propagate streamlines throughout the white matter. 24 major white matter tracts were identified as originally described by Yeatman and colleagues (2012)(79). Each tract was sampled to 100 nodes. Fractional anisotropy (FA), mean diffusivity (MD), and mean kurtosis (MK) were calculated at each node using the Diffusional Kurtosis Model (DKI(80, 81) and axonal water fraction (AWF) was calculated using the White Matter Tract Integrity Model (WMTI(82)).

To assess the extent to which our computational pipeline impacted the results, we also implemented a pipeline that generated the streamlines using anatomically-constrained tractography (54) implemented in MRTRIX3 (83). Tract identification proceeded as above. We then grouped the data based on the behavioral data to make group comparisons across the various tracts. These diffusion features were also used as input to machine learning algorithms to predict reading scores. The results of both tractography pipelines were similar and, ultimately, the DIPY tractography was used.

With neuroimaging data, there are often differences between scanners, which can create variation in data quality and the scale of diffusion properties. This makes comparing images acquired from different sites challenging. To account for between-scanner variation, ComBat harmonization was performed on the tractometry data to correct for any scanner-related variance from the data(68–70). Specifically, we employed the neurocombat_sklearn library in the present analysis to perform ComBat harmonization on the data and remove any scanner or site related variation from the neuroimaging data.

### 2.3 PING

Data used in the preparation of this article were obtained from the Pediatric Imaging, Neurocognition and Genetics (PING) Study database (http://ping.chd.ucsd.edu/). PING was launched in 2009 by the National Institute on Drug Abuse (NIDA) and the Eunice Kennedy Shriver National Institute of Child Health & Human Development (NICHD) as a 2-year project of the American Recovery and Reinvestment Act. The primary goal of PING has been to create a data resource of highly standardized and carefully curated magnetic resonance imaging (MRI) data, comprehensive genotyping data, and developmental and neuropsychological assessments for a large cohort of developing children aged 3 to 20 years. The scientific aim of the project is, by openly sharing these data, to amplify the power and productivity of investigations of healthy and disordered development in children, and to increase understanding of the origins of variation in neurobehavioral phenotypes. For up-to-date information, see http://ping.chd.ucsd.edu/.

The PING dataset consists of neuroimaging and behavioral data from 1,119 subjects between 3 and 20 years old. The imaging protocols and processing steps are outlined in Jernigan et al.(33). In the present analysis, we used the diffusion tensor imaging (DTI) data provided by the authors of the PING study, which was processed using AtlasTrack(37). This processed diffusion data includes the mean FA and MD values in 37 major white matter tracts, as well as the mean FA and MD values for the left hemisphere, right hemisphere, and the whole brain. In addition to the neuroimaging data, the PING dataset also provides behavioral assessments from the NIH Toolbox(84, 85), which includes an Oral Reading Recognition Test for ages 3 and above. We looked at age standardized scores on this reading assessment and classified individuals who scored below one standard deviation from the mean as struggling readers.

### 2.4 ABCD

The ABCD dataset consists of neuroimaging and behavioral data from the first observation of a ten-year longitudinal study. This time point includes data from 11,080 subjects between 8 and 11 years old. The imaging protocols and processing steps are outlined in Casey et al. (34). In the present analysis, we used the diffusion MR data provided in ABCD annual release 4.0. Like the HBN data, the ABCD data were processed using pyAFQ, which resulted in tract profiles for 24 major white matter pathways. We also the diffusion tensor imaging (DTI) data provided NIH data archive, which was processed using AtlasTrack(37). This processed diffusion data includes the mean FA and MD values in 37 major white matter tracts, as well as the mean FA and MD values for the left hemisphere, right hemisphere, and the whole brain In addition to the neuroimaging data, the ABCD dataset also provides the same NIH toolbox Oral Reading Recognition Test (84, 85) found in the PING dataset. We looked at age standardized scores on this reading assessment and classified individuals who scored below one standard deviation from the mean as struggling readers.

### 2.5 HCP-YA

The HCP-YA dataset(73) consists of neuroimaging and behavioral data from 1,200 healthy young adults between the ages of 22 and 35. As with the HBN dataset, we used pyAFQ to perform tractometry on these data. For this analysis, specified parameters that yielded tract profiles for 24 major white matter pathways and four diffusion properties, FA, MD, MK (mean kurtosis), and AWF (axonal water fraction) for each tract. As a behavioral measure, we used the NIH Toolbox Oral Reading Recognition Test to explore the relationship between the white matter and reading skill in this sample.

### 2.6 High-dimensional modeling of the relationship between reading and white matter in HBN, PING, ABCD, and HCP-YA

To generate reading score predictions from diffusion properties of the white matter, we fit a series of gradient-boosted random forest models (XGBoost(48)). For each dataset, we fit three XGBoost models using either demographic information, white matter properties, or both demographic information and white matter properties as predictor variables. For all three datasets, we used age, SES, and geographic location (if available) as predictors. The white matter predictors we used depended on the dataset and the tractometry software used to calculate diffusion properties. For the HBN and HCP-YA datasets, we used FA, MD, AWF, and MK in the 24 tracts identified by pyAFQ, while in the PING dataset we used the mean FA and MD values in the 37 tracts identified by AtlasTrack. Before being entered into the XGBoost models, each predictor variable was demeaned and scaled to unit variance. For each model, hyperparameter tuning was performed using a Bayesian optimization procedure (86) that samples over the hyperparameter space and performs 5-fold cross-validation to identify the best model fit. The final model fits were assessed using a held-out test set that was counterbalanced with the training data to have similar demographic properties.

## 3. Results

### 3.1 A dynamic relationship between changes in the white matter and growth in reading skill

Past longitudinal and intervention studies have suggested that there is not a stable relationship between white matter properties and reading skill but, rather, that diffusion measures of the white matter and reading ability both change dramatically depending on an individual’s educational environment(27, 28, 30). To test the hypothesis that the white matter is part of a dynamic, experience-dependent system that is linked with learning over time, we used a large, longitudinal sample to model the relationship between changes in white matter diffusion properties and growth in reading skills. Here we define reading skills as single word reading and rely on different assessments, depending on the dataset, that largely tap into the same latent construct. These assessments have been used interchangeably across past studies investigating the relationship between white matter and reading skill (Supplementary Table 1). Further, based on past findings, we focused on the left arcuate and the left ILF since these tracts are typically considered to be part of the core reading circuitry(26).

### 3.2 Development of the left arcuate tracks reading development

We first explored the relationship between changes in white matter properties in the left arcuate and changes in reading scores using data from the Pediatric Longitudinal Imaging, Neurocognition, and Genetics study (PLING(36)). In this dataset, diffusion properties of the white matter were calculated using AtlasTrack(37), which provides mean FA and MD (mean diffusivity) for 37 white matter tracts. To explore the relationship between reading and white matter in the PLING sample, we fit a longitudinal linear mixed-effects model to predict mean-centered TOWRE (Test of Reading Word Efficiency) reading scores using time point, initial age, and mean-centered FA in the left arcuate, and mean FA in the left arcuate as fixed-effects and subject as random effects. The inclusion of both mean-centered FA at each time point and overall mean FA allowed us to separate year to year change within an individual from mean FA differences between subjects.

This model revealed significant effects of time point (t(229) =13.98, p < 0.001), and mean-centered FA in the left arcuate (t(299)=2.36, p = 0.019). The effect of mean-centered FA suggests that, within an individual, changes in the diffusion properties of the left arcuate are linked with changes in reading scores across time, even after controlling for age-related increases in FA and overall FA level (Figure 1A). Interestingly, we find no stable relationship between reading skill and white matter properties when we examine the correlation between TOWRE scores and FA in the left arcuate at each time point separately (Supplemental Figure 1). Taken together, these results suggest that within individual changes in diffusion properties of the left arcuate fasciculus track gains in learning over time and call into question the notion of stable individual differences in white matter structure.

**Figure 1:**
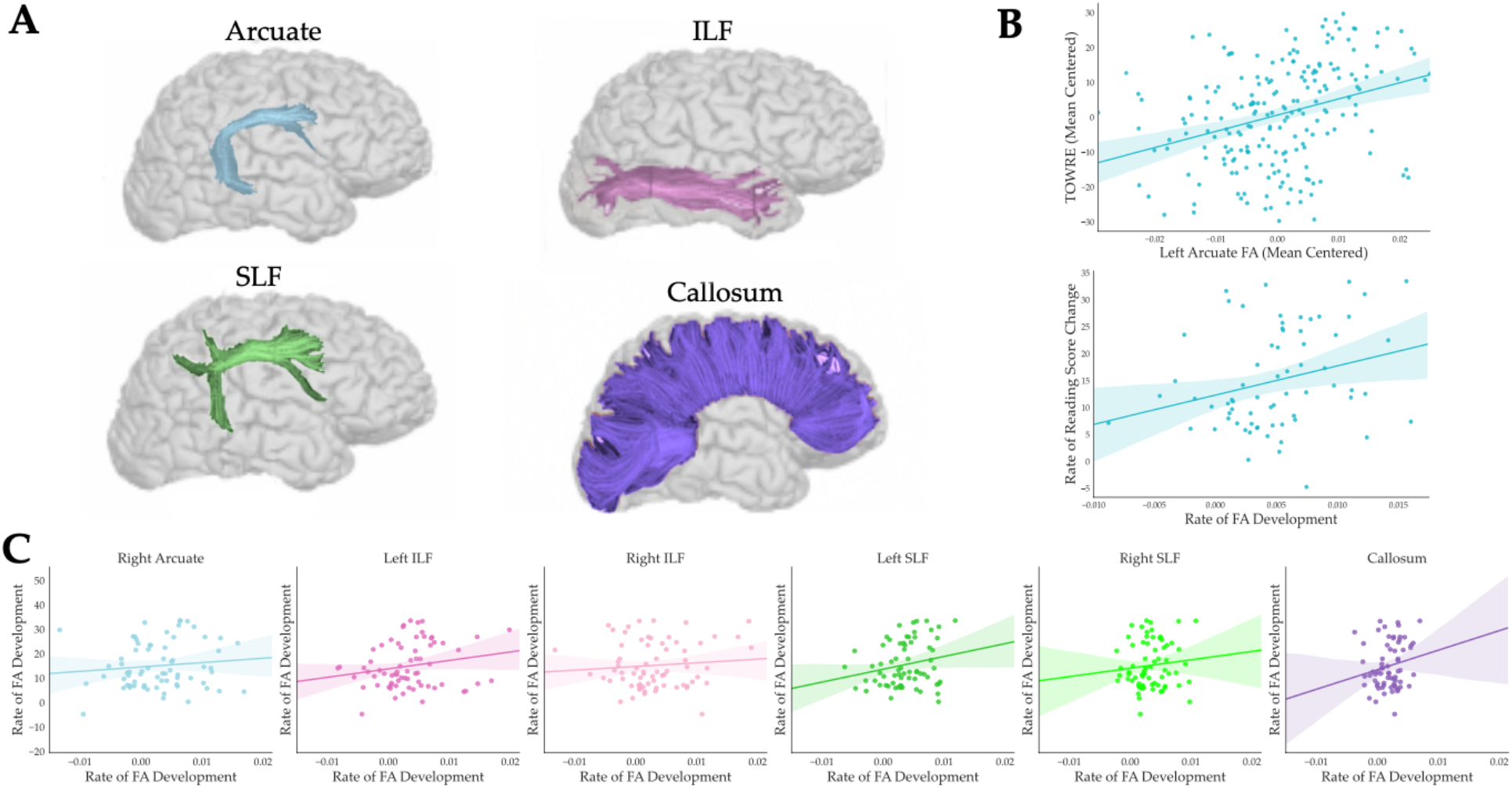
**A**: Rendering of the arcuate, inferior longitudinal fasciculus, superior longitudinal fasciculus, and corpus callosum identified with AtlasTrack. Due to constraints with the PLING dataset, only renderings of the right arcuate, ILF, and SLF are available. **B.** Upper Panel: Mean-centered FA (Fractional Anisotropy) values at each time point for the left arcuate correlate with mean centered reading scores assessed at each time point, illustrating the longitudinal relationship between changes in FA and changes in reading skill. Lower Panel: Relationship between individual rates of FA development in the left arcuate and rate of raw TOWRE (Test of Reading Word Efficiency) reading score change. **C**: Relationship between individual rates of FA development in the right Arcuate, left and right ILF, left and right SLF, and callosum and rate of raw TOWRE reading score change in the longitudinal PLING dataset.

To estimate growth rates for FA, we fit a linear regression model within each individual, predicting each subject’s FA value as a function of age in years (similar to Yeatman et al.(27)). The coefficients of this linear model serve as an estimate of the rate of change in FA over time within a given white matter tract. We applied a similar approach to generate individual estimates of reading score change over time.

We first examined the correlation between the rate of FA development and the rate of raw reading score improvement and found a significant relationship between the two (r = 0.27, adjusted p = 0.02, Figure 1B). Thus, children with more rapid growth in reading abilities also show more rapid white matter development in the left arcuate. We next used the individual growth rates to examine the relationship between the rate of FA development and initial age to see if FA growth rates in the left arcuate differed across developmental stages. This correlation was not significant (r = −0.210, p = 0.08), suggesting that the rate of FA change in the left arcuate is not significantly different from linear over the age range present in the sample.

### 3.3 Reading skills predict future FA development in the left arcuate

Given the relationship between changes in FA in the left arcuate and changes in reading skill, we then tested whether changes in one variable preceded changes in the other or if these changes occurred in parallel. To model the longitudinal interplay between white matter development and reading, we constructed a multi-level vector autoregression model (mlVAR(38, 39)). Within this model, we used the time series of FA measurements in the left arcuate and TOWRE reading scores to assess whether reading scores at a given time point are explained by FA at the previous time point or vice versa. This analysis revealed that FA in the left arcuate at one time point does not predict reading scores at the next time point (t(162) = −0.01, p = 0.971), but that reading skill at one time point does, in fact, predict FA at the following time point (t(162) = 5.44, p < 0.0001).

To compare the beta-coefficients of the model path connecting past reading scores with future FA against the path linking past FA with future reading scores, we conducted a bootstrapped difference test (40). To do so, we generated a bootstrap distribution (n=2,000) of mlVAR coefficients and constructed a 95% confidence interval around the difference between the reading to FA coefficient and the FA to reading coefficient. The interval of the difference between the two coefficients did not include 0 (Bootstrapped 95% CI: [0.728, 0.748]) suggesting that the two coefficients are significantly different. Thus, growth in reading skills from one year to the next predicts future changes in the white matter, whereas developmental changes in the white matter do not predict future gains in reading skill.

The mlVAR model revealed that reading skill at one time point predicts FA in the left arcuate at the next, however, this model does not provide insight into the relationship between the rate of reading development and the rate of FA development in the left arcuate. To better understand this dynamic, we constructed a parallel process latent growth curve model of the longitudinal growth of reading skill, FA, and the relationship between the two. Because the model did not converge properly due to an insufficient number of participants at the fifth time point, only observations from the first four time points were included in this analysis. Figure 2 presents a path diagram illustrating the hypothesized growth trajectories of reading skill and FA in the left arcuate. To assess the impact of change in reading score on change in FA (and vice versa), we generated two models incorporating two different regressors (in place of covariance structures): one predicting the slope of FA from the slope of reading and another predicting the slope of reading from the slope of FA. Based on modeling of individual growth rates presented above, we defined this latent growth curve using linear growth rates for both reading and FA. The overall model fits were acceptable (χ2(22) = 33.41, p=0.056, RMSEA = 0.085, TLI = 0.973). Additionally, the regression predicting FA slope from reading slope was significant (z=1.956, p=0.05), whereas the regression predicting reading slope from FA slope was not (z=-1.311, p=0.190). Taken together, the results from these models suggest that, while both FA and reading skill develop over time, the rate of individual reading gains predicts the rate of future FA development, while the rate of FA development does not predict the rate of reading gains.

**Figure 2:**
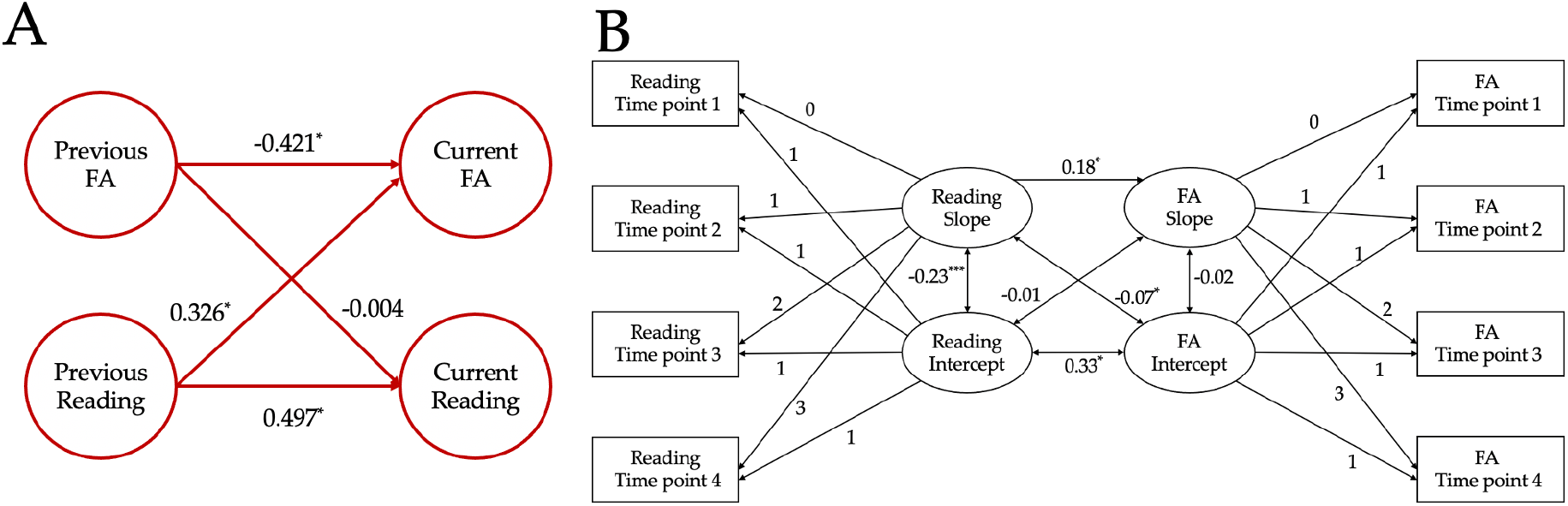
**A.** Path diagram illustrating the mlVAR model capturing the temporal dynamics between the development of reading and FA in the left arcuate over time. The values represent the beta-weights associated with each path within the model. **B.** Path diagram outlining the parallel process latent growth curve model capturing the interplay between longitudinal growth in reading and FA development in the left arcuate. Rectangles represent observed reading or FA values at a given time point and ovals represent latent slope and intercept variables. Unidirectional arrows represent regressions within the model and the adjacent numbers signify the associated beta-coefficients. Bidirectional arrows represent covariance structures and the adjacent numbers signify the covariance values.

### 3.4 Rate of FA development in the right arcuate and left ILF are not related to changes in reading

To test whether the same longitudinal relationships existed between reading and additional white matter tracts, we first constructed the same longitudinal mixed-effects model in the right arcuate, a tract that has not been implicated in the literature as a core component of the reading circuitry. This model revealed significant main effects of initial age (t(66) = 7.15, p < 0.00001) and time point (t(165) = 14.92, p < 0.00001) but no effect of FA in the right arcuate (t(159)=1.104, p = 0.2712). We also conducted the same longitudinal mixed-effect analysis using FA in the left ILF since past findings have also identified the left ILF as part of the reading circuitry. Again this model revealed significant main effects of initial age (t(72)=7.46, p < 0.00001) and time point (t(186)=15.16, p < 0.00001). However, we did not observe a significant main effect of FA in the left ILF (t(198)=0.85, p=0.400). These main effects suggest that neither FA in the right arcuate nor the left ILF significantly relate to changes in reading skills over time. There was also no significant relationship between FA in right arcuate or ILF and reading scores when the time points were examined separately.

### 3.5 No static relationships between white matter properties and reading ability

After establishing the dynamic, longitudinal relationship between white matter development and reading development, we then attempted to replicate past findings from the literature which have suggested that static differences in the white matter explain individual differences in reading skill. Using the HBN dataset(32), one of the most diverse large-scale pediatric samples to date, we compared the white matter diffusion properties of struggling readers (again using a typical cutoff for dyslexia) and a control group.

The HBN data was processed with QSIPrep(41), rigorous quality control was applied to each participant’s data(42), and tractometry was performed with pyAFQ(43). After identifying 24 major white matter tracts, pyAFQ samples each tract to 100 nodes and provides various diffusion metrics for each node (see Methods for overview). The following node-wise group comparisons focused on the left arcuate, left ILF, left SLF, and corpus callosum, which have been most consistently implicated in past studies examining the relationship between the white matter and reading skill(26). Based on past cross-sectional findings, the expectation is that children with low reading scores will show reduced FA values in the left arcuate, left ILF and left SLF compared to the control group and increased FA in the posterior callosum. Additionally, we analyzed 16 control tracts to examine anatomical specificity in the large and diverse HBN sample (Supplemental Figure 2).

An initial group comparison using minimal quality control and not controlling for age or SES revealed reduced FA in areas of the left cingulum cingulate, right cerebral spinal tract, and the callosal motor fibers in struggling readers (corrected p < 0.05, Supplemental Figure 3), but not in left arcuate, left ILF, or left SLF (all corrected p>0.05). After filtering for quality control (See Methods), performing ComBat harmonization on the tract profile data to remove the effects of the scan site, and controlling for various confounds, a group comparison at each node failed to reveal any significant differences in FA (Figure 3, all adjusted p >0.05). Figure 3 shows the effect of reading group at each node after controlling for age, SES and geographic location. There were no nodes with significant group differences. Since the HBN data contains a broad distribution of ages, we repeated this group comparison separately for three age bins as roughly found in the literature: 5-9 years old (27, 28, 30, 44), 10-15 years old (29, 45, 46), 14-21 years old (47). As with the full sample, we found no group differences across any of the age ranges (Supplementary Figure 7).

**Figure 3:**
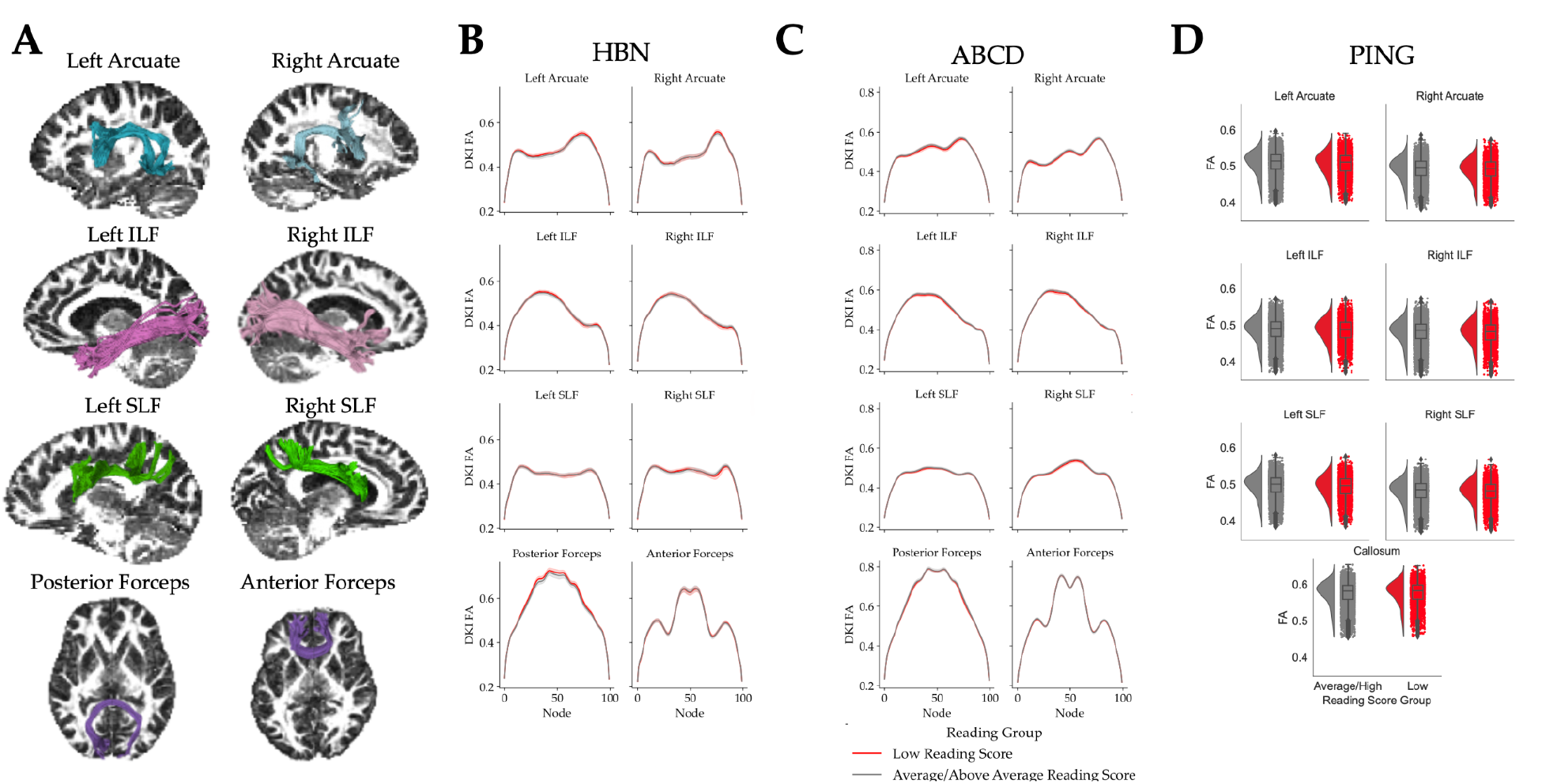
**A:** Rendering of the eight tracts of interest identified with pyAFQ overlaid on the T1-weighted image of the corresponding hemisphere. Note: the posterior and anterior forceps are included as separate tracts though both are part of the corpus callosum. **B**: FA (Fractional Anisotropy) profiles for the core reading circuitry in the HBN data derived by the default pyAFQ pipeline for reading groups after controlling for age and harmonizing on scanner site. The shaded area around each line represents one standard error. **C**: FA profiles for the core reading circuitry in the ABCD. **D**: Rain cloud plots showing the distributions of mean FA values for the core reading circuitry by reading group in the PING data set. There are no significant differences in the diffusion properties between reading score groups across the two datasets. There is a negligible effect of reading group on FA values across all seven tracts.

To validate the surprising lack of group differences observed in the HBN dataset, we also explored the relationship between white matter and reading in the PING dataset(32–34). As with the PLING data, tractometry was performed on the PING data using AtlasTrack (as opposed to pyAFQ; see Methods for overview). Therefore, we used mean FA and MD for 37 white matter tracts in this analysis. In these data, we first looked to see if we could detect differences in the mean FA across the major white matter tracts identified by AtlasTrack. After controlling for age and parental income, we again did not find any significant differences in mean FA between the struggling readers and control groups (Figure 3, all p > 0.05).

After observing no group differences in the HBN and PING datasets, we turned to the ABCD dataset(32–34), the largest ongoing developmental neuroimaging study. Similar to the HBN dataset, tractometry was performed using pyAFQ. We constructed a series of regression models to explore the extent to which reading skills predict FA in the core reading circuitry. After controlling for demographic factors, such as parental income, neighborhood deprivation, and school achievement, the addition of reading group to the model had a negligible effect on the amount of variance explained between the two models (all ΔR^2^ < 0.003, all all Cohen’s f^2^ < 0.003, Figure 3).

### 3.6 Diffusion properties fail to predict reading scores above demographics in three large-scale datasets

After failing to observe any significant differences between individuals with low and high reading scores, we explored the idea that white matter differences related to reading skill are not localized to a single tract but, rather, form a distributed network that might have a nonlinear relationship to reading ability. To test this hypothesis, we fit a series of gradient-boosted random forest models (XGBoost(48)) to determine how well diffusion properties from the entire white matter, not just the reading circuitry, serve to predict reading scores. For this particular analysis, we prioritized predictive accuracy (as opposed to interpretability)(49). We chose the XGBoost algorithm because of its exceptional performance across a wide range of machine learning applications(50–52) and its ability to capture complex, non-linear relationships that would be missed by a linear regression model.

Our first model contained only age, SES, and geographic location as predictor variables; the second model only contained diffusion properties (FA, MD, AWF, MK) from all 24 tracts delineated in the dataset; the third model contained both demographic variables and white matter properties as predictors. Each predictor was first demeaned and then scaled to unit variance before being entered into the XGBoost models.

Cross-validated model R^2^ showed that the demographics-only and white-matter-only models explain roughly the same amount of variance in reading scores(Figure 4). The combination of white matter features and demographic information as predictors did not improve model fit above the models with only demographic information or only white matter features. Thus, as expected, differences in demographics (e.g., SES) did explain some variance in reading scores(53–55). The model with only diffusion properties did predict roughly 5% of the variance in reading scores (Table 1). However, white matter properties did not explain additional variance in reading ability above and beyond demographics indicating that there was not a specific relationship between reading skill and the white matter, beyond what is predicted by sociodemographic factors (Table 1). Furthermore, when we divided the sample into low, medium, and high SES bins and applied the same modeling pipeline, neither demographic features nor white matter properties explained any variance in reading skill. We then turned to a different measure of reading ability, the Test of Word Reading Efficiency (TOWRE), to test whether our WIAT composite score was effectively tapping into the construct of reading skill. Similar to our first set of models, diffusion properties of the white matter did not explain additional variance in reading skill above and beyond demographic factors (Supplemental Figure 4).

**Figure 4:**
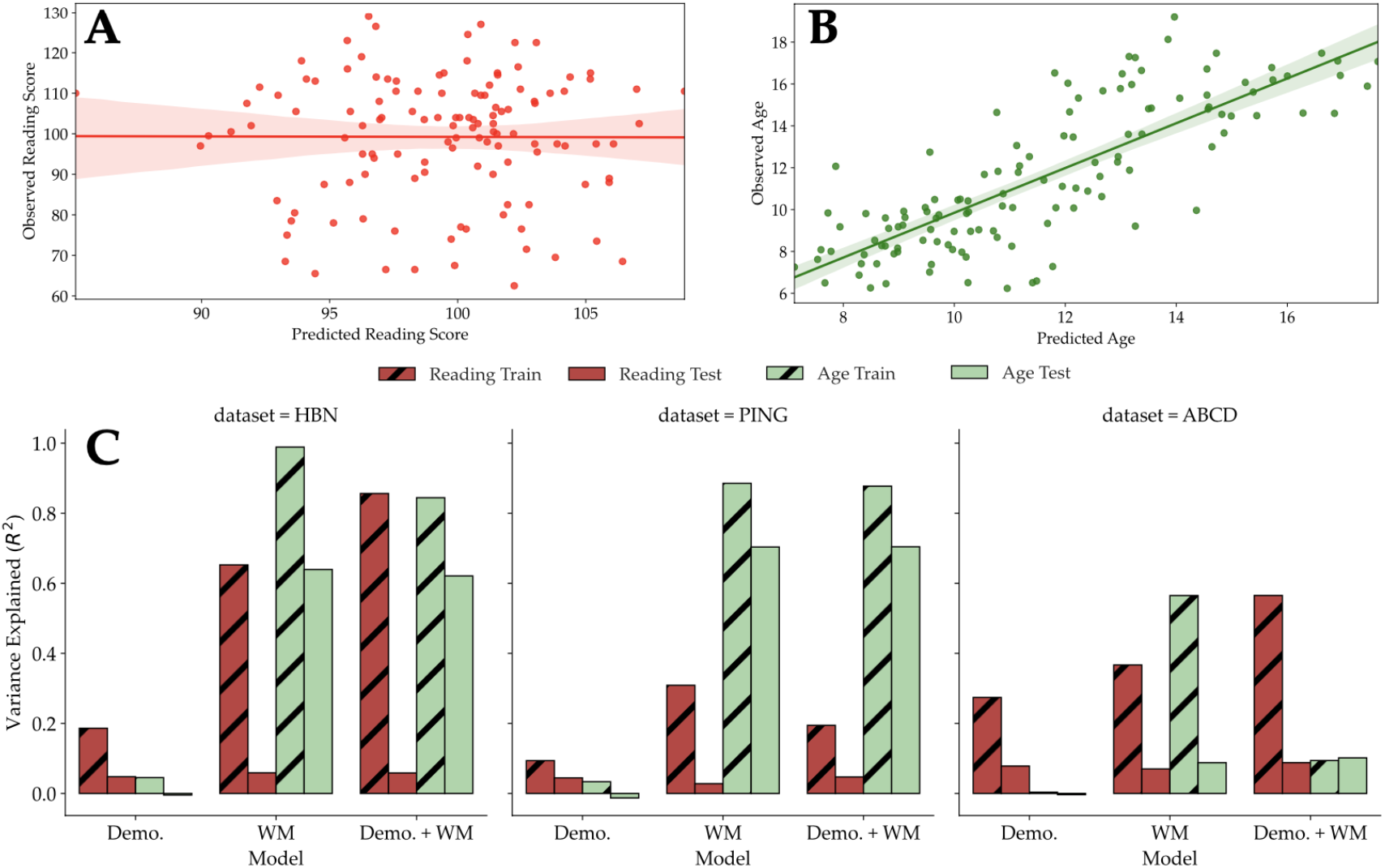
**A.** Predicted vs observed reading scores of the test data generated by the XGBoost model trained on white matter features from the HBN data (R^2^=0.059). **B.** Predicted vs. observed age from XGBoost model trained on white matter features from HBN data (R^2^=0.64). **C.** Train and test R^2^ scores for XGBoost models predicting reading (red) and age (green) in the HBN (left panel) and PING (center panel) and ABCD (right panel) datasets.

**Table 1:** Cross-validated R^2^ test scores for reading score predictions made from XGBoost models trained on white matter properties. The diffusion properties used to train the models vary due to differences in the tractography software used on the various data sets.

| Dataset | Diffusion Predictors | Number of Prediction Features | Cross-validated $R^2$ Score (Test Set) |
| --- | --- | --- | --- |
| HBN - Full Sample | FA, MD, MK, AWF | 9,600 | 0.059 |
| HBN - Full Sample | FA | 2,400 | 0.061 |
| HBN - Full Sample | MD | 2,400 | 0.045 |
| HBN - Full Sample | MK | 2,400 | -0.102 |
| HBN - Full Sample | AWF | 2,400 | -0.024 |
| HBN - Full Sample | Mean FA | 24 | -0.025 |
| HBN - Full Sample | Mean MD | 24 | -0.013 |
| HBN - Full Sample | Mean MK | 24 | -0.034 |
| HBN - Full Sample | Mean AWF | 24 | 0.012 |
| HBN - Low SES | FA, MD, MK, AWF | 9,600 | -0.057 |
| HBN - Medium SES | FA, MD, MK, AWF | 9,600 | -0.178 |
| HBN - High SES | FA, MD, MK, AWF | 9,600 | -0.075 |
| HBN - Site CBIC | FA, MD, MK, AWF | 9,600 | -0.192 |
| HBN - Site CUNY | FA, MD, MK, AWF | 9,600 | -1.004 |
| HBN - Site RU | FA, MD, MK, AWF | 9,600 | 0.053 |
| HBN - Homogenous subset | FA, MD, MK, AWF | 9,600 | 0.240 |
| HCP-YA - Top/Bottom 100 Readers | FA, MD, MK, AWF | 9,600 | 0.079 |
| PING - Homogenous subset | Mean FA, Mean MD | 37 | -0.044 |

To further examine the surprising finding that individual differences in the white matter do not, in fact, predict reading abilities (above and beyond demographic factors), we reprocessed the data from scratch using a different approach to tractography. Specifically, we re-calculated tractometry properties using tractography data generated by QSIPrep(53) using anatomically constrained tractography(54) (ACT; see Methods) to see if the lack of reading ability prediction stemmed from our choice of tractography parameters during the initial processing pipeline. These new tractometry properties again revealed group differences in the tract profiles before quality control, but these differences disappeared when we applied more rigorous quality control and harmonized the data across scanner sites (Supplemental Figure 5). We then conducted the same XGBoost modeling approach with these data and again found that white matter properties did not predict reading scores above and beyond demographic information (Supplemental Figure 6).

Finally, we used the outputs of AtlasTrack present in the PING and ABCD datasets to train a series of XGBoost models to predict reading scores from either demographic factors, diffusion properties of the white matter, or both demographic factors and white matter. The models trained on the PING data showed that demographic features, namely socioeconomic status, predicted reading scores (cross-validated R^2^= 0.040) and that the addition white matter features to the model did not increase predictive power (cross-validated R^2^=0.045, t(9)=0.383, p = 0.36, Figure 4C). The models trained on the ABCD data also revealed that demographic features explained roughly 10% of the variance in reading scores (cross-validated R^2^= 0.101), and white matter features did not improve predictive power of the model (cross-validated R^2^=0.078).

### 3.7 Maturational differences in white matter properties predict age

To ensure that the lack of replication of the cross-sectional effects did not reflect poor data quality or modeling errors, we sought to benchmark our models against a finding with a well-established effect size. The largest and most consistent effect in the diffusion MRI literature is that FA increases and MD decreases with age(10, 11, 55, 56). Based on this, we would expect to be able to explain a significant portion of the variance in age in the present sample using diffusion data. To evaluate the signal quality of the diffusion data, we fit models to predict each subject’s age using the same XGBoost models. As expected, the demographics-only model failed to predict age above chance confirming that there was not a sampling bias. The white matter model, on the other hand, predicted 64% of the variance in age in the HBN dataset and 70% of the variance in age in the PING dataset (Figure 4C). These predictions are consistent with the state-of-the-art in “brain age” calculations (57) confirming, first, that the diffusion MRI data was sufficiently high quality to accurately index white matter maturation and, second, that our modeling approach was able to accurately capture these effects.

In contrast to the HBN and PING datasets, white matter properties only predicted about 7% of the variance in age in the ABCD dataset. However, all the participants in the ABCD sample are within two years of age of each other and, therefore, there is not much maturational variation in the white matter properties for the model to learn.

### 3.8 Reading score - white matter relationships are sample-dependent

The present analysis leveraged the largest diffusion MRI sample to date to explore the relationship between white matter and reading abilities. Contrary to the compendium of small, single-observation studies, this large sample did not reveal any stable predictors of reading ability in the white matter. One potential explanation for this discrepancy is that previous studies were picking up on real effects that reflected specific characteristics of the small, homogenous samples that do not generalize to the population at large. The HBN dataset is an extremely neurodiverse sample of children from a variety of demographic backgrounds, with 87% of participants having at least one diagnosis and 53% of participants having more than one diagnosis.

To test the hypothesis that seemingly conflicting results across past studies stem from specific features of the participant recruitment strategies, we tested whether we could predict reading scores using a small, homogeneous sub-sample of the entire HBN dataset that excluded participants based on strict quality control measures and co-occurring diagnoses. We began by selecting all the participants between the ages of 8 and 12 years old from one site (CBIC) and excluding any participants diagnosed with autism, anxiety, or ADHD, as these diagnoses have been shown to correlate with reading skills in different ways(58–60). Within this limited subsample, we then identified the 40 top and bottom readers to create groups that differed dramatically in terms of reading ability. The resulting, single-site sample was skewed towards high SES participants, as is the case with many single-laboratory studies. We split this subsample into a training set of 60 subjects and a test set of 20 subjects. We then trained an XGBoost classifier on the train set using 5-fold cross-validation before predicting reading level (high or low) in the test set. The classifier trained on white matter properties had a classification accuracy of 0.70 in the held-out test sample. In addition to this classification model, we trained an XGBoost model to predict reading scores in this same sub-sample, to facilitate comparison across our other modeling pipelines. The model trained on white matter data to predict reading scores in this sub-sample predicts roughly 24% of the variance in reading scores in the test set (Table 1). Further, when we thresholded the reading score predictions made by our regression model using the maximum score for the low reading group as a cutoff, we found that the model had a classification accuracy of 0.80 in the test sample.

Next, we examined the relationship between white matter and reading ability in a large-scale, dataset consisting of healthy, college-aged young adults (HCP). The original HCP recruitment screened for many different developmental, neurological and psychiatric disorders and reflects a more homogenous sample than, for example, the HBN data(61, 62) and is substantially less diverse than the US population. Using an XGBoost model, we found that diffusion properties of the white matter were able to predict 7.9% of the variance in reading scores in the top and bottom 100 readers, in line with other predictions of cognitive ability made using the HCP dataset(63). Thus, in these two cases representing samples that were a) more demographically homogeneous, b) differed substantially in terms of reading ability and, arguably c) are more similar to a typical sample collected by a single university research group, white matter properties did predict individual differences in reading ability.

However, using the same approach in the PING dataset, we were unable to predict reading scores from diffusion properties. Similar to the HBN data, we generated this subsample by taking the 40 highest vs lowest scoring readers between the ages of 8 and 12 years old. We again trained XGBoost models on the data from this subset and were unable to predict reading scores from the average diffusion properties of the white matter (Table 1). The lack of prediction in the PING subset suggests that mean diffusion properties may not be suitable for generating cross-sectional behavioral predictions, even in homogenous samples.

## 4. Discussion and Conclusions

We explored two hypotheses surrounding the relationship between white matter and reading skills: a) that white matter and reading skill are dynamically linked over time and change with factors such as an individual’s educational environment and b) that white matter properties act as static biomarkers that predict differences in reading abilities. To test the first hypothesis, we examined a five-year longitudinal dataset, which revealed that individual growth rates in the left arcuate are linked with growth in reading scores over time. Further modeling revealed that gains in reading predict future changes in FA in the left arcuate fasciculus. These models suggest that individual differences in learning are related to white matter development rather than white matter differences serving as constraints on the learning process.

To explore the second hypothesis, we analyzed three large-scale cross-sectional datasets (HBN: n=777, ages: 5-21; PING: n=1,119, ages: 3-20; ABCD: n=11,080, ages: 8-11) and found no stable relationship between white matter diffusion properties and reading scores. To capitalize on the strengths of both explanatory and predictive modeling(49), we conducted group comparisons to explore the specific white matter properties underlying differences in reading skill and also leveraged high-dimensional models to predict reading scores from white matter properties. Across all three of these datasets, univariate group comparisons revealed no differences between poor and skilled readers and XGBoost models were unable to predict reading scores from diffusion features (after controlling for SES). However, analysis of a large-scale adult dataset (HCP-YA: n = 1,200) and a subset of the HBN data revealed that diffusion properties of the white matter serve to predict reading scores in more homogenous samples, suggesting that group differences in the white matter between typical and struggling readers may depend on the makeup of the sample. All together, these results suggest that properties of the white matter may not necessarily serve as static traits that differentiate individuals but, rather, that white matter and reading skill may be part of a dynamically linked system that changes over time.

The longitudinal dynamics between reading skill and white matter properties observed in the PLING dataset serve to highlight the role that an individual’s environment may play in driving the development of brain-behavior relationships. In these data, not only did changes in reading skill track changes in FA in the left arcuate, but the rate of individual reading gains also predicted increases in FA in the left arcuate, suggesting gains in reading precede, and potentially influence, changes in the white matter. Other longitudinal studies have shown that skilled readers demonstrate positive rates of FA development in the left arcuate and ILF, whereas poor readers demonstrate shallower developmental slopes in the same tracts (27, 28), but the present study is the first to examine temporal precedence of longitudinal relationships in the reading circuitry. Future work should expand these analyses to additional white matter tracts to better understand the longitudinal relationship between white matter and learning throughout the entire brain, not just within the reading circuitry.

Nevertheless, questions remain as to how growth opportunities within an individual’s learning environment relate to changes in both academic skills and brain properties. In the PLING dataset, only three participants had an initial standardized TOWRE composite score below 85 making it impossible to study the differences in growth trajectories between struggling and typical readers. Additionally, the present sample includes a broad range of ages, and we therefore cannot assess how these longitudinal dynamics change across different stages of development. Future longitudinal and intervention studies will be critical for understanding these developmental dynamics and working out causal relationships between environmental factors, learning, and white matter plasticity.

The longitudinal interplay between white matter growth and reading gains raises questions about the static link between white matter structure and reading skills. Many studies have found differences in the white matter properties of dyslexic versus typical readers based on single observations of small samples(17, 23). However, the present analysis of the HBN, PING, and ABCD datasets (totaling nearly 13,000 participants) failed to reveal any meaningful group differences in the white matter properties of struggling and typical readers. Furthermore, state-of-the art machine learning models were unable to predict reading scores from white matter properties above and beyond demographic factors. This observation is in line with recent work showing that the effect sizes of many brain-behavior relationships are much smaller (or non-existent) than estimates from small samples(31).

These results are also generally in line with those reported by Meisler and Gabrieli(35) which did not find any significant relationship between FA and reading skills in a subset of the HBN dataset. Though the authors observed a relationship between FA and phonemic decoding in the right SLF and the left ICP in participants above age 9, their analysis focused on a different measure of reading skill (TOWRE) and relied on a different analytic pipeline (TractSeg(64)), which may explain the subtle differences between their findings and the present results. Regardless of the differences between these independent analyses of the HBN dataset, both sets of results suggest that the relationship between the white matter and reading skill may not necessarily be reducible to static, individual differences in the diffusion properties of the white matter. White matter properties are highly plastic and changes in the diffusion signal have been detected on the timescale of hours(8) and months(29, 65) after a learning experience. Therefore, it is possible that these datasets serve as a snapshot of a dynamic, experience-dependent system in flux and do not capture the longitudinal relationships between brain, behavior, and environment.

Based on this interpretation, however, one might expect to observe a more stable relationship between reading and white matter properties in adults. Although, when we divide the HBN sample into distinct age bins, we are unable to observe a relationship between reading skill and white matter properties, even in the oldest age range (Supplemental Figures 7 and 8), in the HCP-YA adult sample, we do observe a link between white matter properties and reading skill. Together, these results suggest that environmental and/or developmental factors may dynamically influence the relationship between reading and white matter over the course of development and lead to detectable group differences in adulthood. However, it could also be the case that there exist dynamic traits moderating the relationship between white matter and literacy that obscure group differences over the course of development but lead to detectable differences in adulthood. Future longitudinal studies will be necessary to better understand the dynamics between white matter plasticity and literacy, especially considering that white matter pathways continue to develop throughout childhood and into adulthood (56).

The lack of meaningful group differences in the HBN, PING, and ABCD samples do not necessarily serve to dismiss past findings linking properties of the white matter to reading skill but, rather, present new challenges for the field consider in order to reconcile the seemingly conflicting findings from large-scale data sets, small single-lab studies, and meta-analyses. First, in the present analyses, we rely on traditional diffusion MRI measurements, namely FA. It could be the case that novel diffusion measurements might serve to capture static relationships between the white matter and literacy. Second, as highlighted in recent work using large-scale fMRI datasets, the effect size of various brain-behavior relationships decreases as a function of sample size and makeup (31). Moreover, models that are trained on a biased demographic group often do not generalize to more diverse samples (66). It is possible that sampling procedures impact the relationship between diffusion properties of the white matter and academic skills and may explain the seemingly contradictory results between the present study and past findings.

One of the strengths of these large-scale datasets, especially the HBN sample, is that they include participants from a wide range of geographic and demographic backgrounds, representing diversity that is rarely present in the samples collected by a single lab. Thus, these large-scale datasets provide a new opportunity to explore brain-behavior relationships while controlling for the robust relationship between clinical and socioeconomic factors and academic outcomes. For example, the HBN sample is also extremely neurodiverse: nearly all the individuals in the dataset have some sort of clinical diagnosis. In the final sample used in our analysis, only 35 of 777 participants (4.5%) had no diagnosis whatsoever. Some of these cooccurring diagnoses can impact reading scores and white matter properties and it could be the case that these multiple, overlapping clinical diagnoses may obscure potential brain-behavior relationships. For instance, many individuals diagnosed with autism have been shown to have hyperlexia (58), whereas ADHD or anxiety diagnoses have been linked to lower reading scores (59, 60). On the other hand, past findings from single laboratories that demonstrate a relationship between white matter properties and reading skill are typically based on a single observation of smaller, demographically homogeneous samples, usually recruited from the community surrounding a university. Additionally, these samples often exclude individuals with certain diagnoses. Inconsistent sampling procedures may explain why some studies report increased FA in struggling readers(45, 46) and others report reduced FA in struggling readers(28, 29, 44).

Interestingly, when we generate a small (n=80) homogenous subsample of the HBN data, excluding subjects with autism, ADHD, or anxiety diagnoses, we find that high-dimensional models trained on diffusion properties of the white matter do predict some variance in reading scores. However, it should also be noted that this subset of the data is highly skewed in terms of socioeconomic status, with most of the participants coming from upper SES households. This finding parallels the results observed in the HCP-YA data, which also revealed that properties of the white matter served to predict reading scores in a large, relatively homogenous sample of adults. The recruitment procedures in many single-lab studies may lead to more discernible relationships between individual differences in white matter properties and reading skill but these results might not generalize beyond the sample. In fact, for individuals with high SES backgrounds, genetic factors have been shown to exert more of an influence on FA than those from lower SES backgrounds(67), suggesting that in homogenous, high SES samples, genetic factors may influence the relationship between white matter and reading skill in a manner that may not replicate in more diverse study populations.

These findings also raise questions surrounding the relationship between SES (and other environmental factors more broadly), literacy, and white matter development. In the present analysis, the models trained exclusively on white matter features did predict some variance in reading skills. However, this variance largely overlapped with the variance explained by demographic factors, given that the models trained on both white matter and SES did not serve to explain additional variance in reading abilities and white matter did not predict reading skill whatsoever in models trained on data from distinct SES bins. Previous studies relying on smaller, more homogenous participant populations might have been picking up on this indirect relationship. Untangling the complex relationships between environmental factors such as SES, brain development, and academic achievement is a complex issue (66) and researchers leveraging large-scale, single-observation datasets need to carefully consider how to incorporate covariates, such as clinical diagnoses and SES, into their analyses.

It also bears mention that these large-scale datasets generally consist of data collected using multiple scanners, often made by different manufacturers. Subtle differences across scanners may introduce an additional source of variability to the data despite efforts to harmonize scan sequences across sites (32–34). We employed state-of-the-art approaches to harmonize the neuroimaging data across scanner sites (68–70) and reduce the impact of scanner site on the variability of the diffusion measures. However, when we generated predictive models from each scan site separately, only data from one site predicted any variance in reading scores. Thus, it is possible that differences in scanners and pulse sequences may obscure subtle differences in the white matter that do relate to reading skill.

As the field of developmental cognitive neuroscience moves into the era of Big Data with diverse groups of participants, longitudinal measurements will be of particular importance to answer questions surrounding the relationship between the brain, academic environment, and learning. Past meta-analyses have failed to identify reliable neurological profiles for individuals labeled as “learning disabled”(25, 71), suggesting that a more individualized approach to studying learning may be important, and densely sampled functional neuroimaging studies have found that an individual’s neurobiology is often distinct from group-level patterns (72). A single snapshot of a dynamic system can be misleading and longitudinal studies are critical for capturing the within-individual interplay between environmental factors, brain development and learning.

Historically, the field of developmental cognitive neuroscience has relied on crosssectional samples based on a single observation of each participant to draw connections between brain properties and cognitive skills such as reading. A brief review of the literature reveals that there are at least 27 studies on the neuroanatomical underpinnings of reading skill and of these, only 3 studies consist of more than 2 observations per participant. Furthermore, of the five large-scale, publicly-funded, developmental datasets that we are aware of, only 1 includes observations at least four different time points. Unfortunately, a dynamic system cannot be studied with cross-sectional data - at least four time points are necessary to resolve the dynamic interactions that we discovered in the current work. The present findings provide strong evidence for the dynamic nature of brain-behavior associations and suggest that the developmental cognitive neuroscience should prioritize longitudinal, within-subjects designs. Longitudinal studies examining brain-behavior relationships in tandem with information about an individual’s educational environment will lead to better understanding of how learning opportunities influence brain development and open the door for refined, developmentally appropriate learning interventions and pedagogical strategies.

## Acknowledgements

This manuscript was prepared using a limited access dataset obtained from the Child Mind Institute Biobank, Healthy Brain Network. This manuscript reflects the views of the authors and does not necessarily reflect the opinions or views of the Child Mind Institute.

Additional data were provided [in part] by the Human Connectome Project, WU-Minn Consortium (Principal Investigators: David Van Essen and Kamil Ugurbil; 1U54MH091657) funded by the 16 NIH Institutes and Centers that support the NIH Blueprint for Neuroscience Research; and by the McDonnell Center for Systems Neuroscience at Washington University.

Further data used for this study were obtained from the Pediatric Imaging, Neurocognition and Genetics Study (PING) repository (https://www.nitrc.org/projects/ping), a publicly shared data resource comprising standardized assessments of behavioral, neuroimaging and genetic variables in typically developing children, adolescents and young adults (Jernigan et al., 2016). As such, the investigators within PING (T. L. Jernigan, A. M. Dale, L. Chang, N. Akshoomoff, C. McCabe, E. Newman, T. M. Ernst, P. Van Zijl, J. M. Kuperman, S. S. Murray, C. S. Bloss, N. J. Schork, W. Thompson, H. Bartsch, D. G. Amaral, E. R. Sowell, W. E. Kaufmann, P. Van Zijl, S. Mostofsky, B. J. Casey, B. Rosen, T. Kenet, J. A. Frazier, D. N. Kennedy, and J. R. Gruen) contributed to the design and implementation of PING and/or provided data but did not participate in analysis or writing of this report. Instructions on how to access the data through the NIMH Data Archive (NDA) are provided (https://www.nitrc.org/plugins/mwiki/index.php/ping:MainPage).

Data used in the preparation of this article were obtained from the Adolescent Brain Cognitive Development^SM^ (ABCD) Study (https://abcdstudy.org), held in the NIMH Data Archive (NDA). This is a multisite, longitudinal study designed to recruit more than 10,000 children age 9-10 and follow them over 10 years into early adulthood. The ABCD Study^®^ is supported by the National Institutes of Health and additional federal partners under award numbers U01DA041048, U01DA050989, U01DA051016, U01DA041022, U01DA051018, U01DA051037, U01DA050987, U01DA041174, U01DA041106, U01DA041117, U01DA041028, U01DA041134, U01DA050988, U01DA051039, U01DA041156, U01DA041025, U01DA041120, U01DA051038, U01DA041148, U01DA041093, U01DA041089, U24DA041123, U24DA041147. A full list of supporters is available at https://abcdstudy.org/federal-partners.html. A listing of participating sites and a complete listing of the study investigators can be found at https://abcdstudy.org/consortium_members/. ABCD consortium investigators designed and implemented the study and/or provided data but did not necessarily participate in the analysis or writing of this report. This manuscript reflects the views of the authors and may not reflect the opinions or views of the NIH or ABCD consortium investigators. The ABCD data repository grows and changes over time. The ABCD data used in this report came from http://dx.doi.org/10.15154/1523041.

This work was funded by Eunice Kennedy Shriver National Institute of Child Health and Human Development grants R01HD095861 and P50HD052120 to JDY, RF1MH121868 to AR and JDY and R01EB027585-02 to AR and Eleftherios Garyfallidis. ARH, AR and MN were also additionally supported by the Gordon & Betty Moore Foundation and the Alfred P. Sloan Foundation, through grants to the University of Washington eScience Institute Data Science Environment. Some compute resources were provided through Amazon Web Services and a grant from Microsoft Azure through the University of Washington eScience Institute

## Supplemental Figures

**Supplemental 1:**
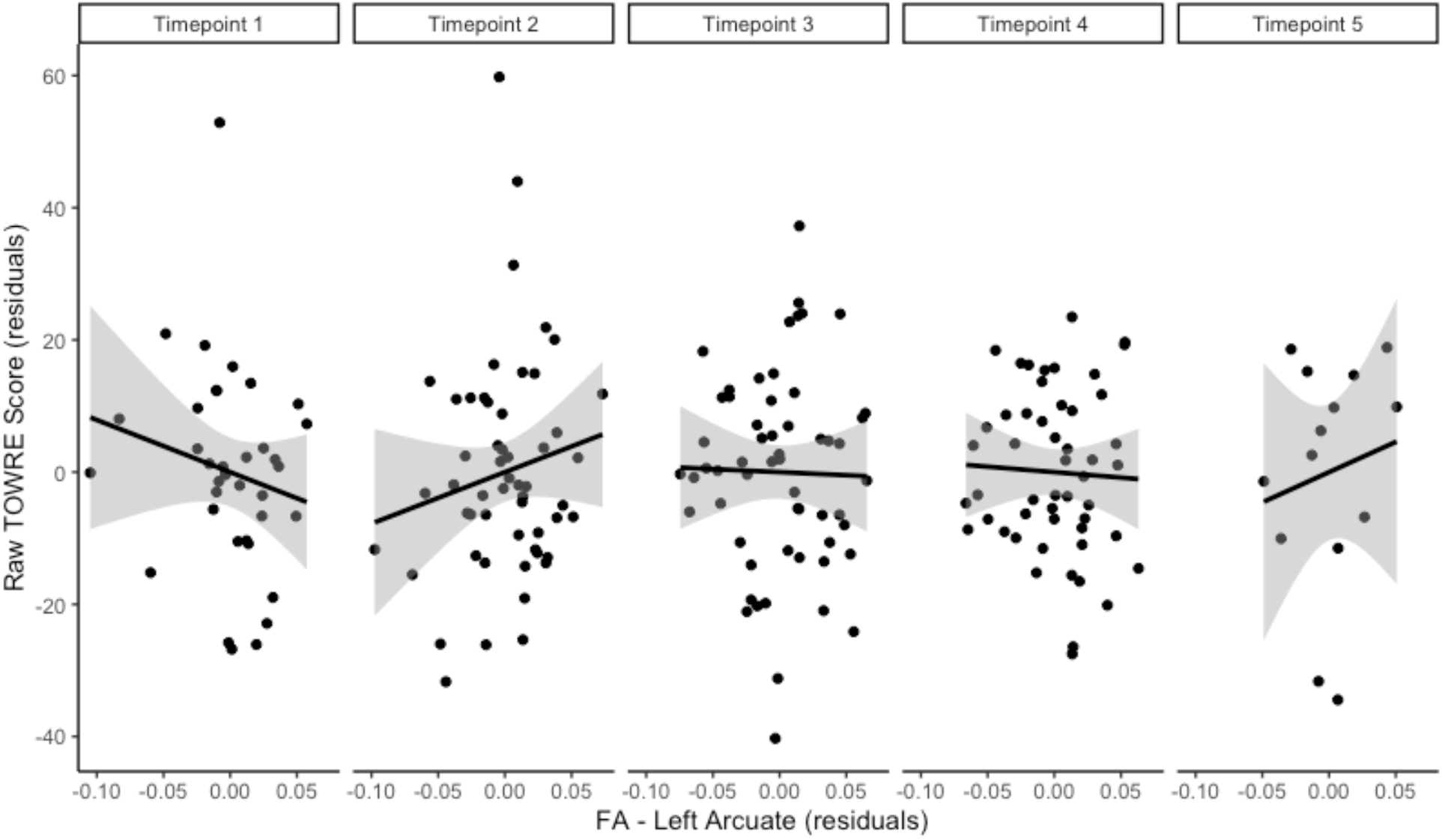
Relationship between age-residualized FA (fractional anisotropy) in the left arcuate and age-residualized raw TOWRE (Test of Reading Word Efficiency) score at each time point in the PLING data.

**Supplemental 2:**
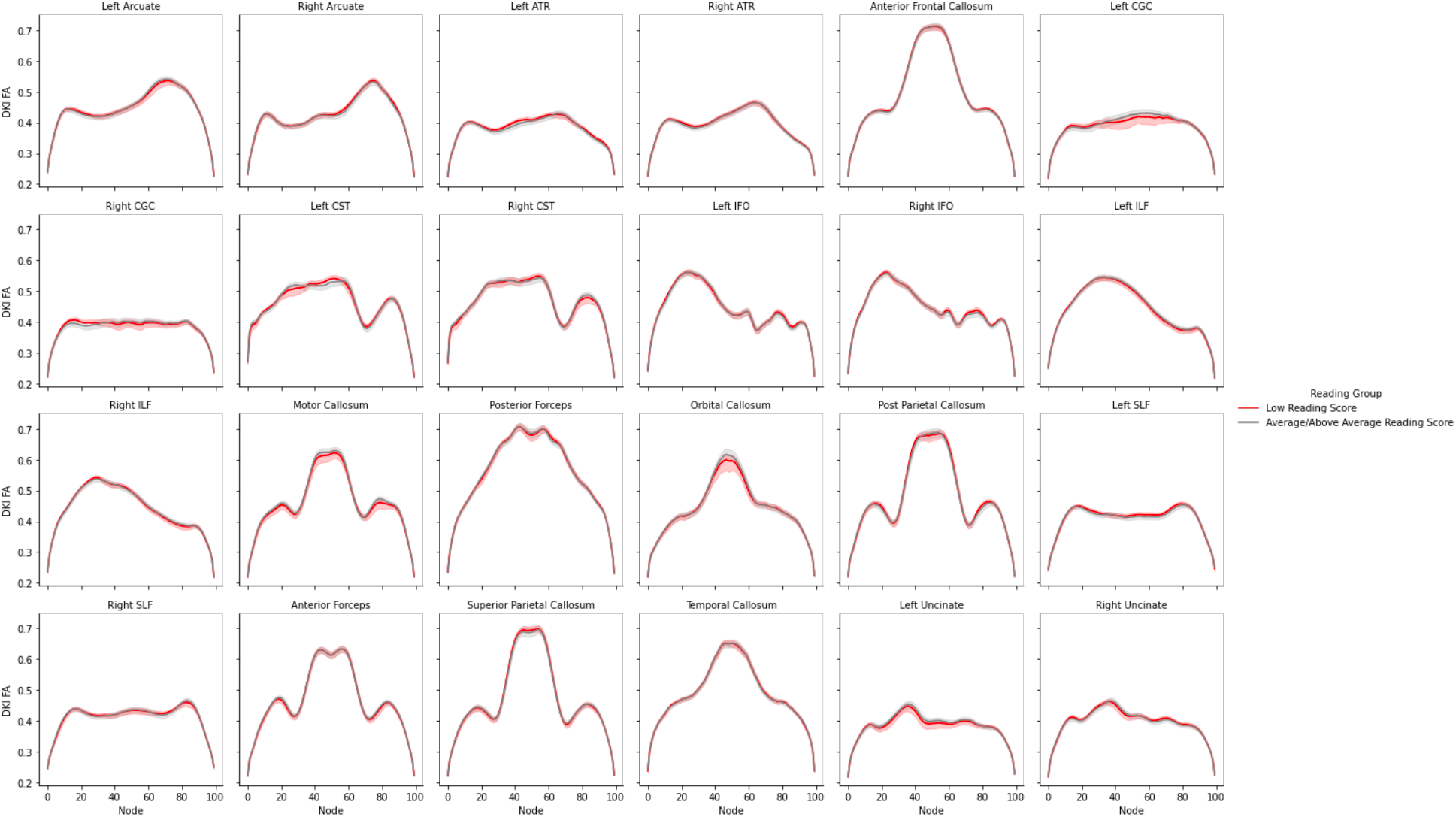
FA profiles for the 24 white matter tracts identified by pyAFQ pipelines on the full HBN sample. The red line represents poor readers and gray lines represent typical readers. After controlling for age and correcting for multiple comparisons, there are no significant differences between the two groups across all 24 white matter tracts.

**Supplemental 3:**
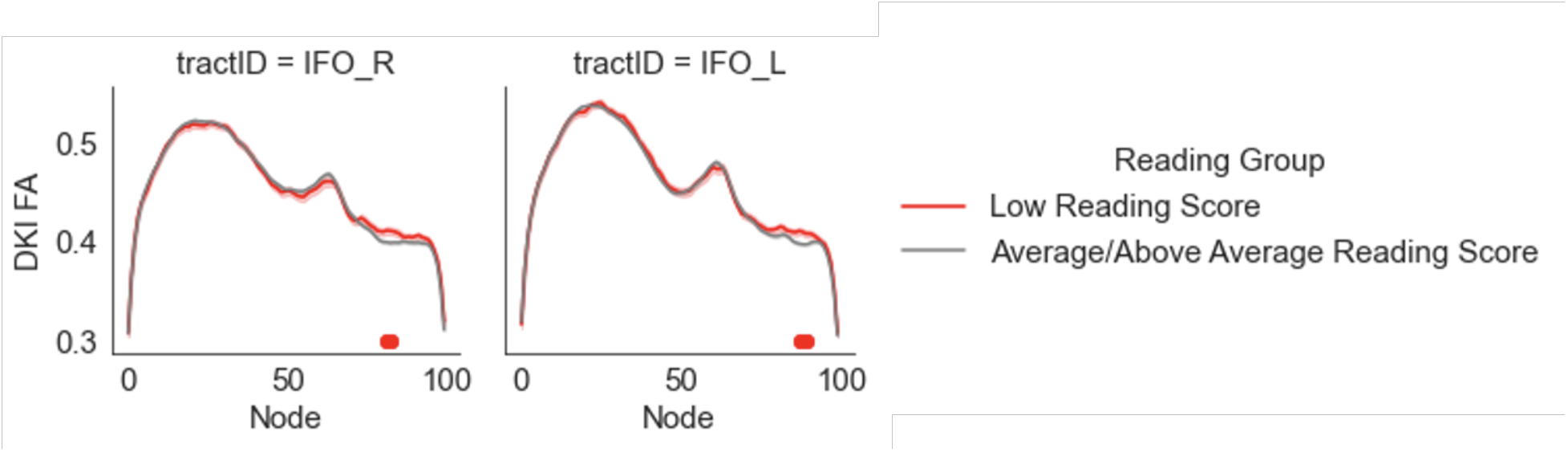
Group differences in naive comparison of reading score groups in a pyAFQ pipeline using anatomically constrained tractography to calculate diffusion properties in the HBN dataset. Average/Above Average readers (gray) demonstrate reduced FA in parts of the left and right IFO. Red dots indicate nodes where the difference in FA is significantly different (adjusted p < 0.05)

**Supplemental 4:**
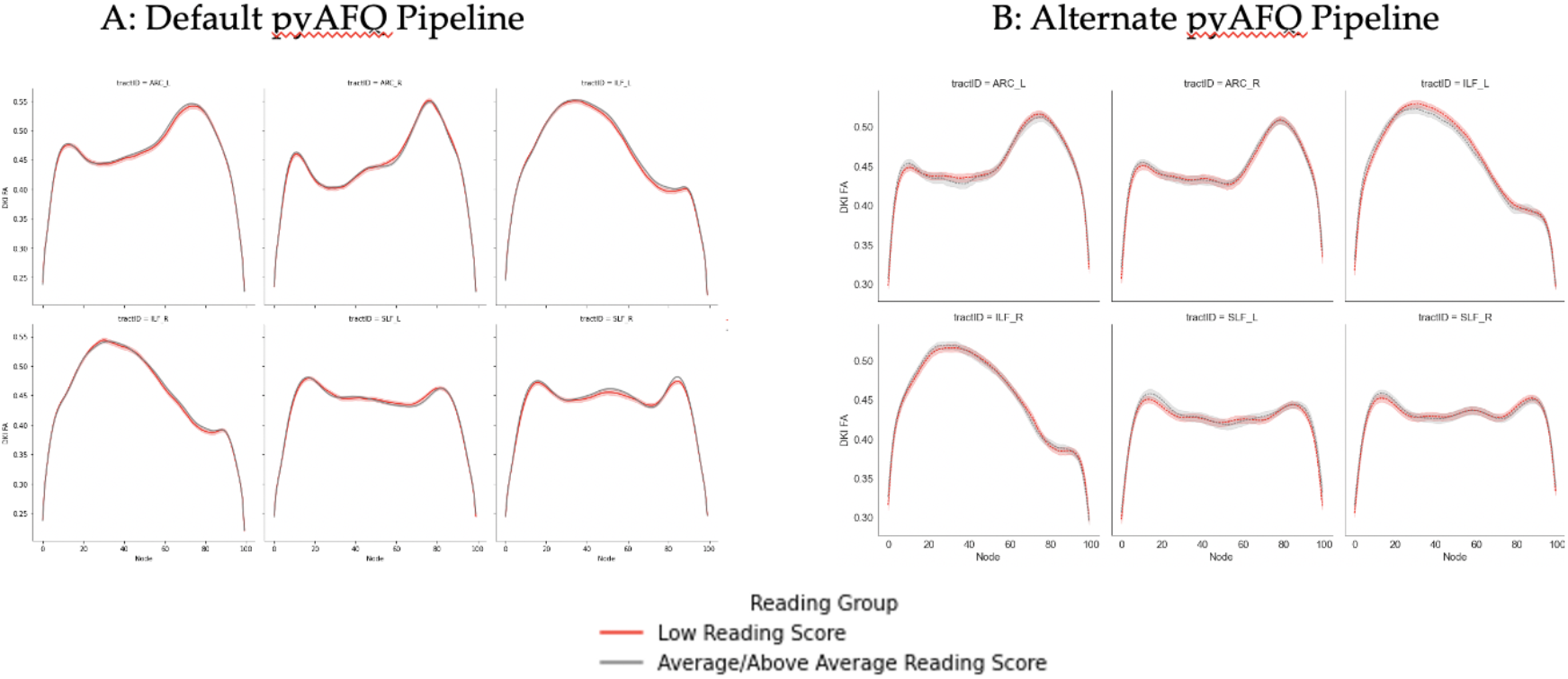
Tract profiles for the left and right arcuate fasciculus, ILF, and SLF as derived from two different pyAFQ pipelines on the full HBN sample. The default pipeline (left) uses probabilistic tractography whereas the right pipeline uses anatomically constrained tractography implemented in MRTRIX3. Neither analytic pipeline reveals significant group differences in the tract profiles of struggling and typical readers.

**Supplemental 5:**
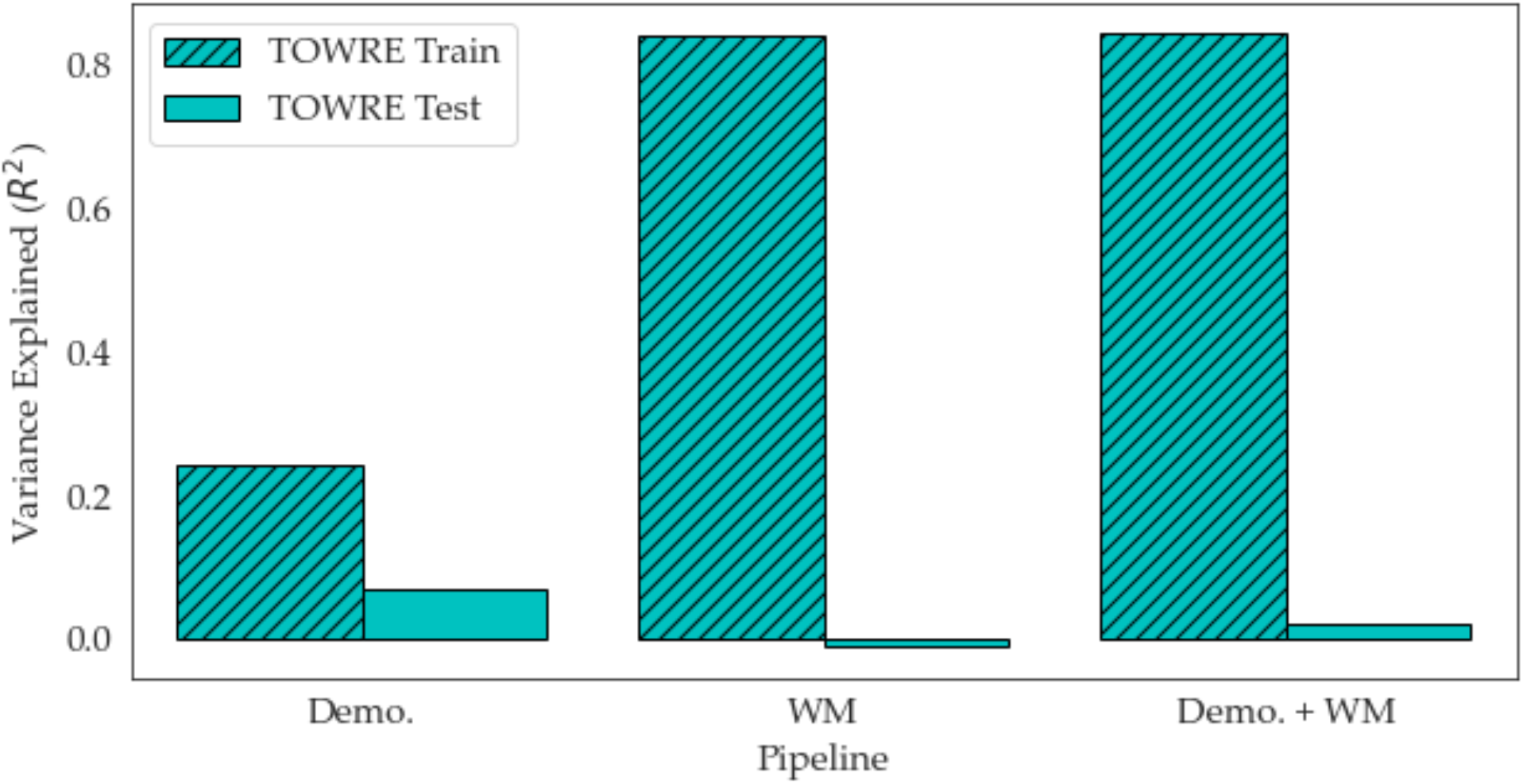
Train and test R^2^ values for the models predicting TOWRE reading skill in the HBN dataset using demographic and white matter features generated by pyAFQ.

**Supplemental 6:**
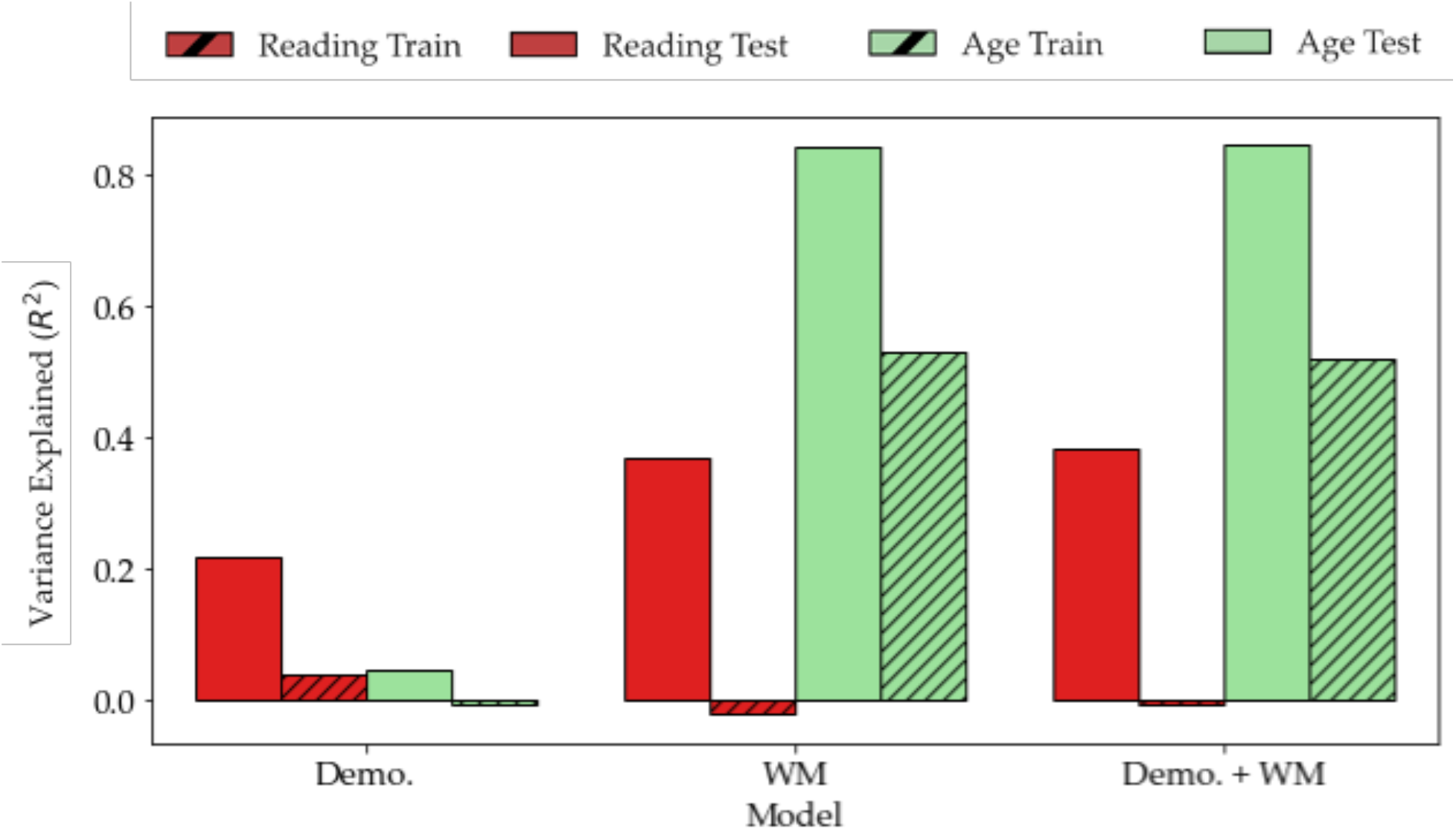
Train and test R^2^ values for the models predicting reading and age in the HBN dataset using demographic and white matter features generated from an alternate pyAFQ pipeline.

**Supplemental 7:**
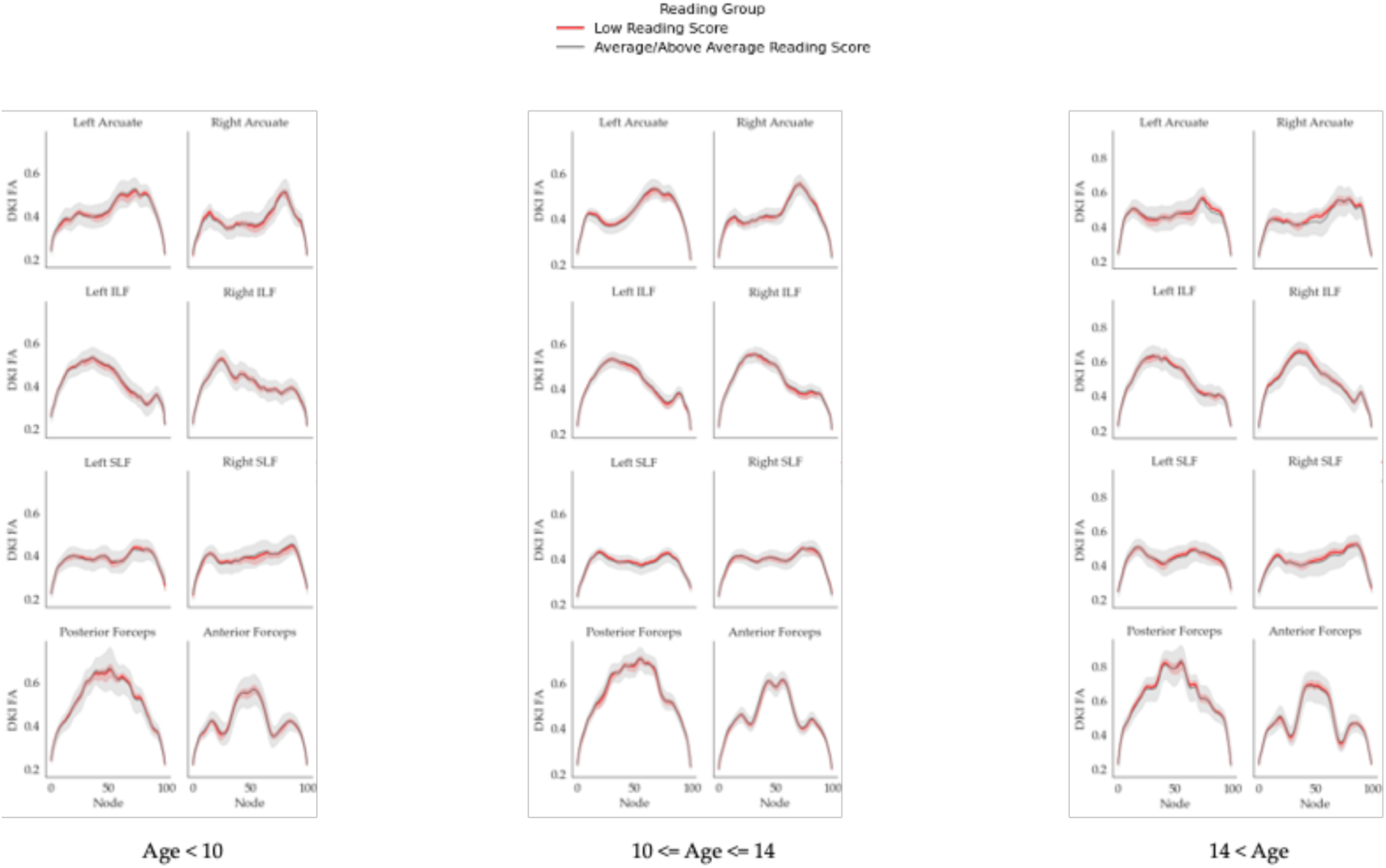
Tract profiles for the left and right arcuate fasciculus, ILF, and SLF for three different age groups (left: age<10; center: 10<=age<=14; right: 14<age) within the HBN sample. The red line represents poor readers and gray lines represent typical readers in each age bin. After correcting for multiple comparisons, there are no significant differences between the two reading groups across the three age bins.

**Supplemental 8:**
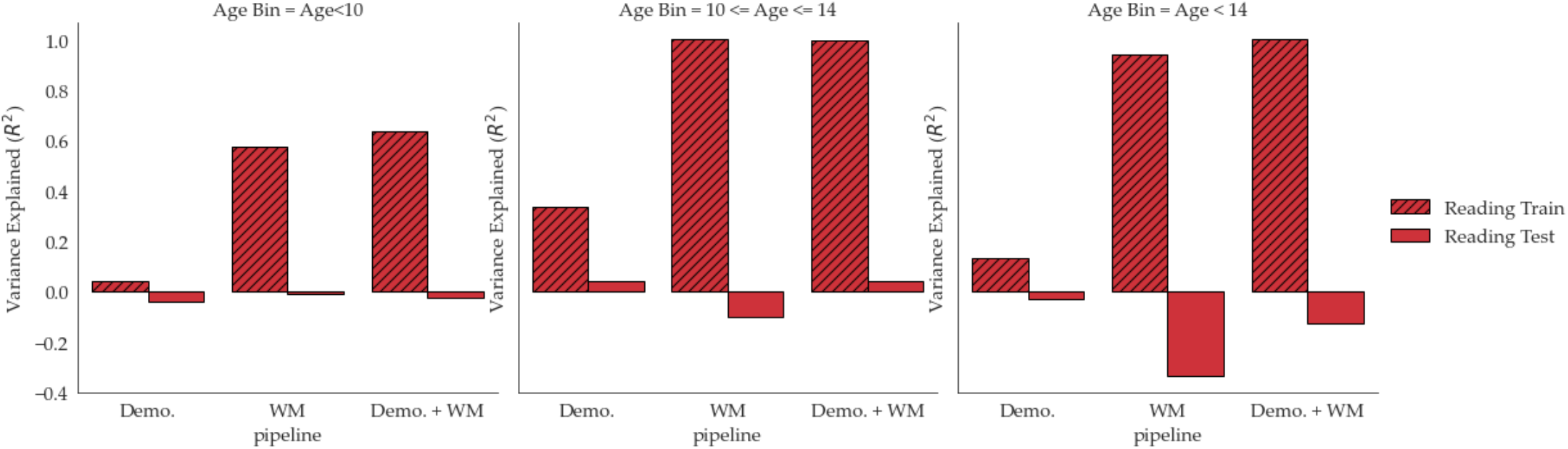
Train and test R^2^ values for the models predicting reading by age group in the HBN dataset using demographic and white matter features generated using pyAFQ. None of the models predicted reading scores above chance.

**Supplementary Table 1:**
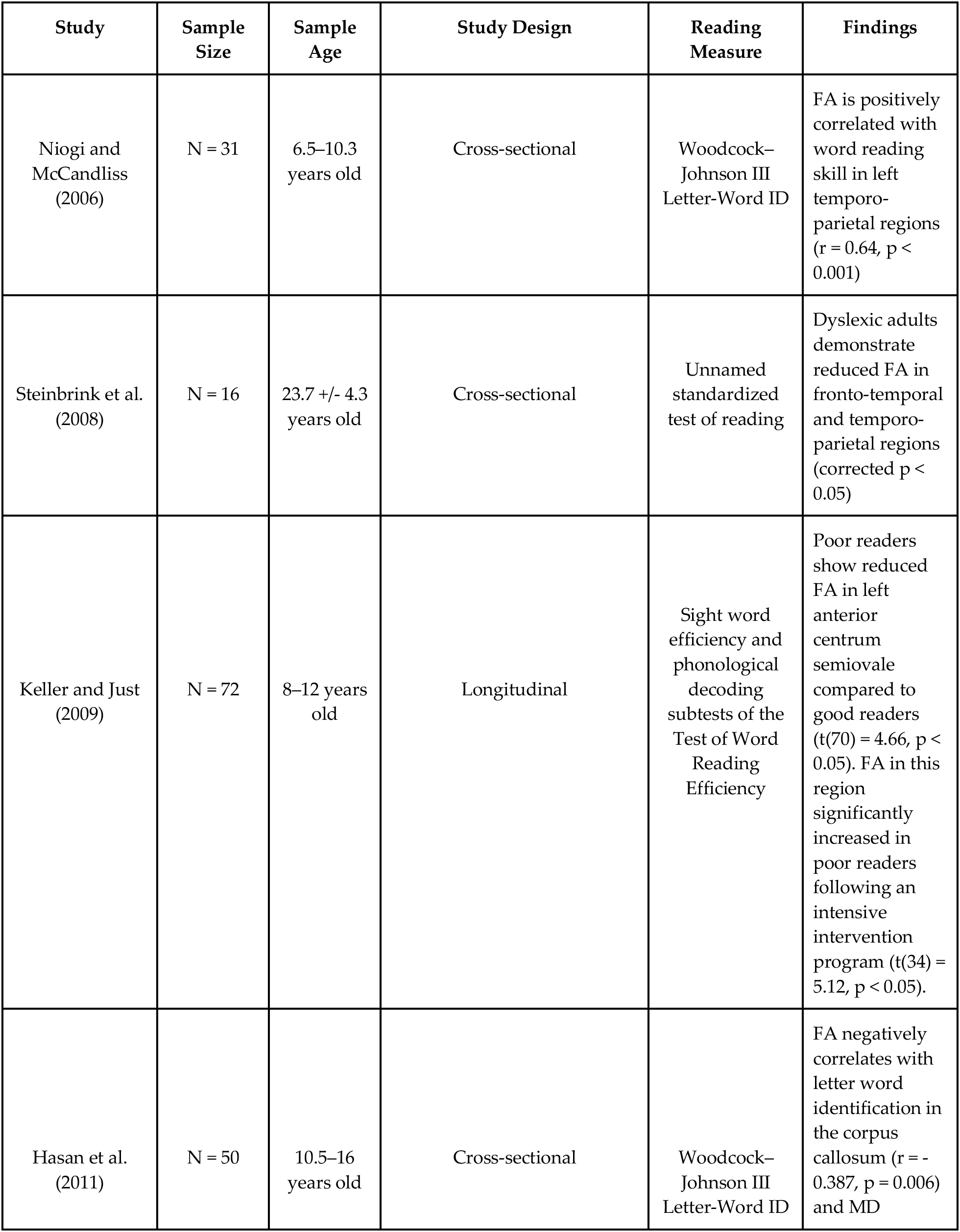
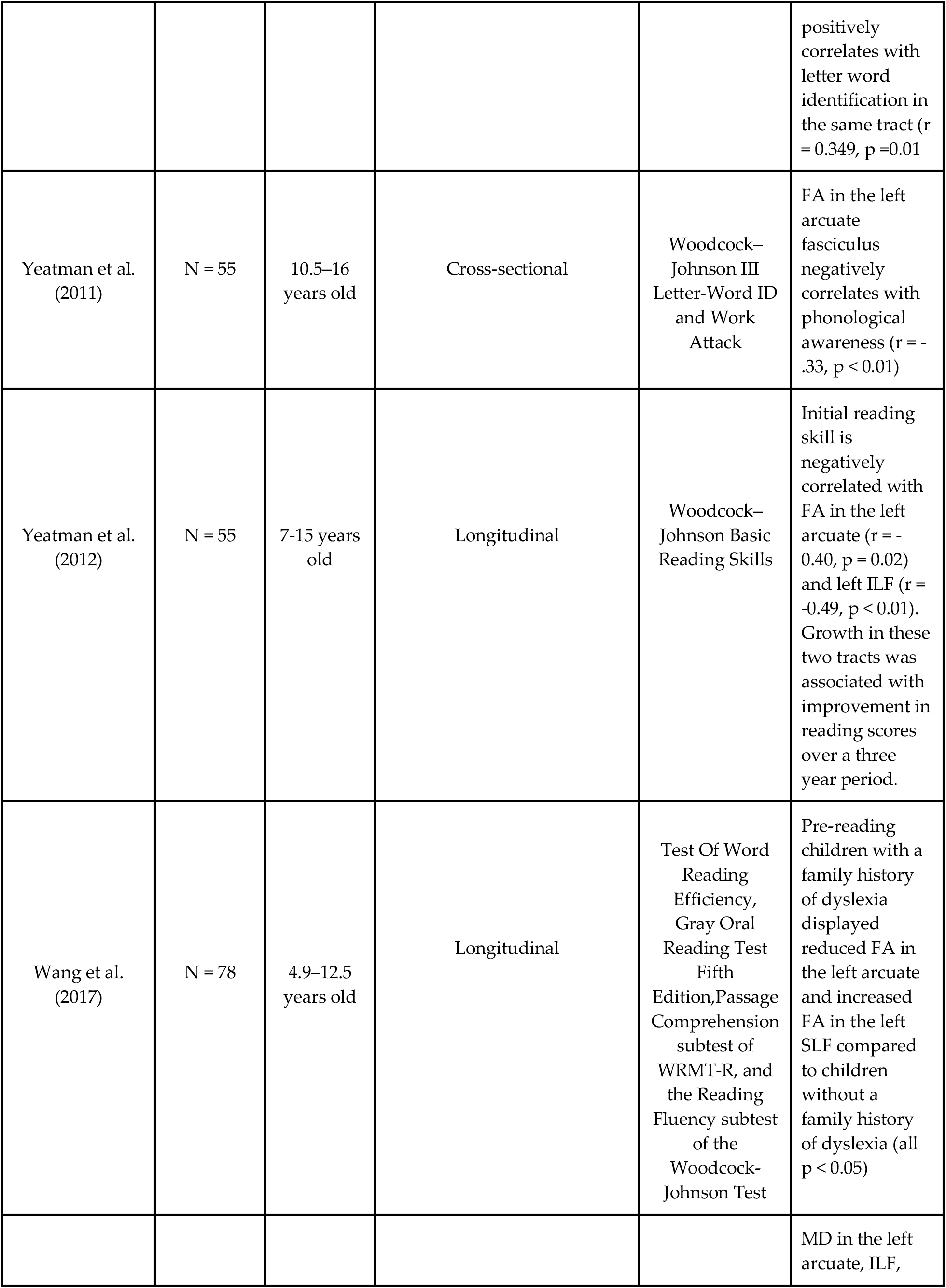
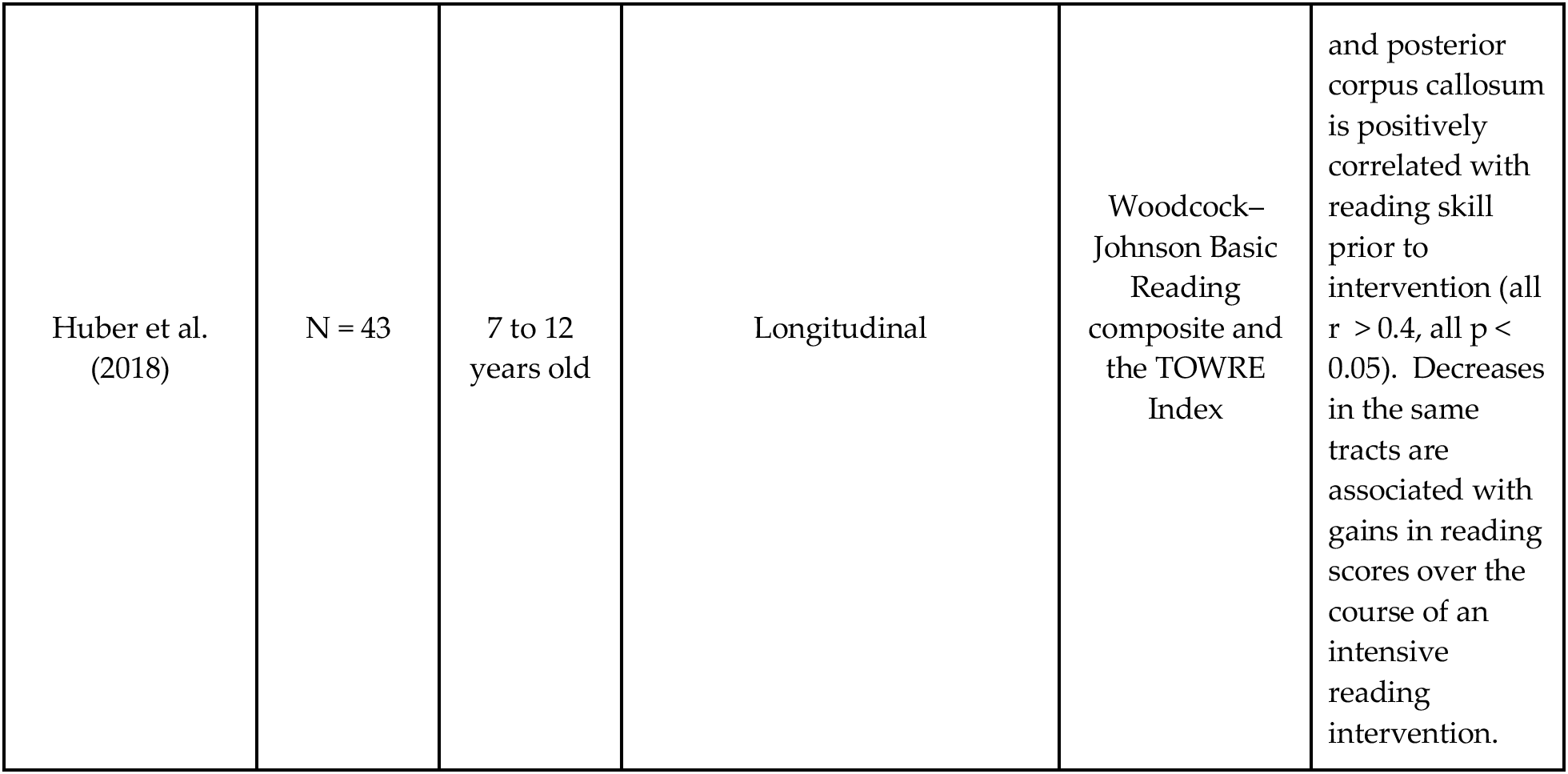
Selective overview of cross-sectional and longitudinal studies investigating the relationship between properties of the white matter and reading skills.

## Supplemental Text

### Prediction of reading scores based on connectivity matrices

Tractometry is one of many approaches to summarizing anatomical properties of the white matter. An alternative approach to analyzing diffusion data is to construct connectivity matrices summarizing the number of streamlines connecting each cortical region. These “structural connectivity matrices” have been used to infer differences in connectivity among cortical regions and also differences in connectivity patterns across subjects. To ensure that the lack of prediction was not specific to our analysis approach (tractometry), we also explored an alternative analysis pipeline that sought to predict reading scores from the structural connectivity matrices generated with the qsiprep reconstruction implementing anatomically constrained tractography based on the multi-shell, multi-tissue model(53, 87). For this approach, we used the same gradient boosted forest pipeline outlined above but, rather than FA and MD values, we used streamline counts from the AAL-VOIs atlas(88) as our white matter predictors. Again, this model did not predict additional variance in reading scores above the demographics only model (Supplemental Table).

| Predictors | Target | Train | Test |
| --- | --- | --- | --- |
| SES + Age | Reading | 0.145 | 0.046 |
| Connectivity | Reading | 0.456 | -0.006 |
| SES + Age + Connectivity | Reading | 0.584 | -0.004 |
| SES | Age | 0.029 | -0.082 |
| Connectivity | Age | 0.846 | 0.587 |

